# A scalar Poincaré map for struggling in *Xenopus* tadpoles

**DOI:** 10.1101/540971

**Authors:** Mark Olenik, Conor Houghton

## Abstract

Short-term synaptic plasticity is widely found in many areas of the central nervous system. In particular, it is believed that synaptic depression can act as a mechanism to allow simple networks to generate a range of different firing patterns. The locomotor circuit of hatchling *Xenopus* tadpoles produces two types of behaviours: swimming and the slower, stronger struggling movement that is associated with rhythmic bends of the whole body. Struggling is accompanied by anti-phase bursts in neurons on each side of the spinal cord and is believed to be governed by a short-term synaptic depression of commissural inhibition. To better understand burst generation in struggling, we study a minimal network of two neurons coupled through depressing inhibitory synapses. Depending on the strength of the synaptic conductance between the two neurons, such a network can produce symmetric *n* – *n* anti-phase bursts, where neurons fire *n* spikes in alternation, with the period of such solutions increasing with the strength of the synaptic conductance. Relying on the timescale disparity in the model, we reduce the eight-dimensional network equations to a fully explicit scalar Poincaré burst map. This map tracks the state of synaptic depression from one burst to the next, and captures the complex bursting dynamics of the network. Fixed points of this map are associated with stable burst solutions of the full network model, and are created through fold bifurcations of maps. We derive conditions that describe period increment bifurcations between stable *n* – *n* and (*n* + 1) – (*n* + 1) bursts, producing a full bifurcation diagram of the burst cycle period. Predictions of the Poincaré map fit excellently with numerical simulations of the full network model, and allow the study of parameter sensitivity for rhythm generation.

## 1. Introduction

Short-term synaptic plasticity may have a role in burst activity in central pattern generators (CPGs). Short-term synaptic depression is commonly found in neuronal networks involved in the generation of rhythmic movements, such as in the pyloric CPG of the spiny lobster (Manor et al., 1997; Rabbah & Nadim, 2007), or in the lumbosacral cord of the chick embryo (Donovan et al., 1998). Synaptic depression modulates the strength of synapses in response to changes to the presynaptic firing frequency. At a high neuronal firing frequency, depression weakens the strength of synapses and therefore reduces the magnitude of the respective postsynaptic response. At low firing frequency, it allows sufficient time for the synapse to recover from depression between spikes, leading to a stronger postsynaptic response. In reciprocal networks, synaptic depression has been shown to act as a “switch”, giving rise to a wide range of network dynamics such as synchronous and multi-stable rhythms, as well as fine tuning the frequency of network oscillations (Nadim & Manor, 2000; Nadim et al., 1999; Bose & Booth, 2011).

A switch between two CPG rhythms that is believed to be guided by synaptic depression can be observed in hatchling *Xenopus* tadpoles. Tadpoles select between two main behaviours in response to different types of skin stimuli. Swimming occurs in response to a brief touch or a single electrical pulse to the trunk (Clarke et al., 1984), and is accompanied by 12 to 25 Hz alternating left and right bending moves (Kahn et al., 1982). Alternatively if held, tadpoles produce struggling, with stronger, slower 2 to 10 Hz bending moves of the whole body (Kahn & Roberts, 1982). The switch from swimming to struggling is associated with a change from a pattern of low-frequency alternating single spikes in swimming (Kahn et al., 1982), to one of alternating high-frequency bursting in struggling (Kahn & Roberts, 1982; Soffe, 1993). Both swimming and struggling are thought to be produced by two overlapping spinal cord CPG networks that share some types of individual neurons. One such shared class of neurons are the inhibitory commissural interneurons (cINs), which have reciprocal depressing inhibitory synapses with neurons on the opposite site of the spinal cord (Li et al., 2007).

Brown (1911) pioneered the idea that synaptic depression acts as a burst termination mechanism in reciprocally inhibitory CPGs involved in rhythm generation of locomotion. Weakening of inhibition as a result of synaptic depression allows the antagonistic side to be released, start firing, and terminate the ongoing burst. Brown’s rhythmogenesis hypothesis has been considered one of a handful of standard mechanisms for generating locomotion rhythms in vertebrates (Reiss, 1962; Perkel & Mulloney, 1974; Friesen, 1994). Brown’s hypothesis is also believed to explain struggling in tadpoles (Li et al., 2007), where the periodic weakening of antagonistic cIN inhibition through short term synaptic depression may act as a burst termination mechanism that produces the antiphase burst rhythm.

Here we study Brown’s rhythmogenesis hypothesis and the mechanisms of burst generation in struggling using a generic half-centre CPG that consists of two identical, tonically active Morris-Lecar (Morris & Lecar, 1981) neurons that are coupled through inhibitory depressing synapses. Each neuron represents a population of cINs on either side of the tadpole spinal cord. Our model reduction is based on the approach previously taken by Bose & Booth (2011), where the dynamics of a similar two-cell reciprocal inhibitory network with depression has been analysed. Bose & Booth’s numerical simulations showed that when the reciprocal synaptic conductance between the two neurons was varied, the model produces symmetric *n* – *n* anti-phase bursts, with stronger synaptic coupling leading to longer bursts (see fig. 3). Bose & Booth used methods from geometric singular perturbation theory to separate the timescales of the fast membrane, and the slow synaptic dynamics of the network and derive one-dimensional conditions for the existence of stable *n* – *n* solutions (for *n* < 3). Most importantly, they showed that the type of pattern largely depends on the slow depression dynamics of the synapses between the two neurons, and can therefore be predicted by knowing the strengths of the synaptic conductances of the two synapses.

In this work we extend the analysis by Bose & Booth (2011) by reducing a two-cell model to a scalar Poincaré map that tracks the evolution of the depression from the beginning of one burst to the beginning of the next burst. Fixed points of our map are associated with stable *n* – *n* burst solutions. Our map construction is motivated by the burst length map of a T-type calcium current, utilised in Matveev et al. (2007), which approximates the anti-phase bursting dynamics of a network of two coupled Morris-Lecar neurons. In contrast to our model, the Matveev et al. network does not contain short-term synaptic depression, and burst termination is instead accomplished through the dynamics of a slow T-type calcium current.

Our Poincaré map replicates the results from numerical simulations of the full two-cell ODE system: Given the strength of maximum conductance between the two neurons, fixed points of our map predict the type and period of *n* – *n* patterns, the switch between burst solutions of different periods, as well as the occurrence of co-existent solutions. We show that fixed points are created via a fold bifurcation of maps, and we derive algebraic conditions that predict the period increment bifurcations that allow the switch between *n* – *n* and (*n* + 1) – (*n* + 1) solutions. Because our map is fully explicit, it lays the framework for studying the effects of any other model parameter on network dynamics without the need to run expensive numerical integrations of the ODEs. Our analysis suggests that understanding the dynamics of the slow depression is sufficient to explain the generation of burst solutions in the tadpole struggling network.

This paper is organised as follows. First, we introduce the network of two neurons, and describe the properties of single cell and synapse dynamics. We use numerical simulations of the network to provide an intuition for the range of possible burst dynamics the system can produce. Next, we state and justify the simplifying assumptions that are necessary for the map construction. Finally, we analytically derive the first return map of the depression variable as well as the conditions that are required for stable solutions to bifurcate between *n* – *n* and (*n* +1) – (*n* + 1) solutions. We end this work with a discussion.

## 2. Model

We consider a pair of identical Morris-Lecar neurons (Morris & Lecar, 1981), with parameters adapted from Bose & Booth (2011). The Morris-Lecar model is a set of two first-order differential equations that describe the membrane dynamics of a spiking neuron. The depolarisation is modelled by an instantaneous calcium current, and the hyperpolarisation by a slow potassium current and a leak current. The membrane potential *v_i_* and potassium activation *w_i_* of neuron *i*(*i, j* = 1, 2) is described by:

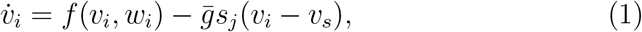

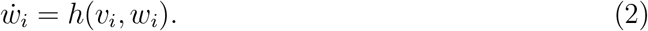

Here *v_s_* is the inhibitory reversal potential, and 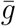 and *s_j_* are the maximal synaptic conductance and the synaptic gating, respectively, constituting the total inhibitory conductance 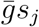 from neuron *j* to neuron *i*. Function *f*(*v_i_*, *w_i_*) describes the membrane currents of a single cell:

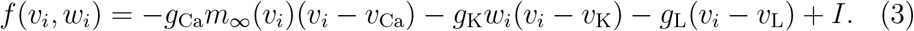

The currents include a constant current *I*, and three ionic currents: an instantaneous calcium current, a potassium current, and a leak current, with respective reversal potentials *v*_Ca_, *v*_K_, and *v*_L_, as well as maximum conductances *g*_Ca_, *g*_K_, and *g*_L_. The function *h*(*v_i_*, *w_i_*) models the kinetics of the potassium gating variable *w_i_*, and is given by

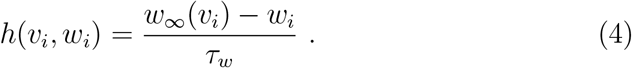

The steady-state activation functions *m*_∞_ and *w*_∞_ as well as the default model parameters are described in Appendix A.

The dynamics of the synaptic interactions between the neurons are governed by a synaptic gating variable *s_i_* and a depression variable *d_i_*:

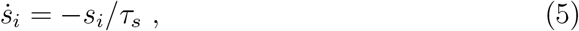

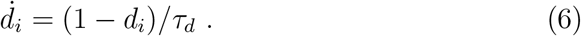

Variable *d_i_* describes a firing rate dependent depletion mechanism that governs the amount of depression acting on the synapse. Our model is agnostic with respect to the exact mechanism of this depletion, be it pre- or post-synaptic. In between spikes *s_i_* decays with time constant *τ_s_*, while *d_i_* recovers with time constant *τ_d_*. Because synaptic depression occurs on a much slower timescale than synaptic inhibition, we assume *τ_d_* ≫ *τ_s_*. At spike time when the voltage *v_i_* crosses a synaptic threshold *v_θ_*, the synaptic variable *s_i_* is reset to the current value of *d_i_*, and *d_i_* is scaled down by the depression strength parameter λ, where 0 < λ < 1. This results in the following discontinuous reset rule on spike time:

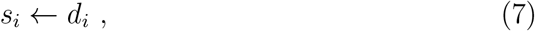

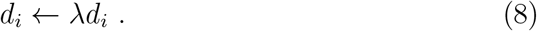

The equations for the depression model were adapted from Bose et al. (2001). These equations are a mathematically tractable simplification of the established phenomenological depression model previously described by Tsodyks & Markram (1997). The original equations in Bose et al. include an exponential decay of *d_i_* throughout the active phase of the action potential, that is 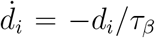 for *v_i_* > *v_θ_*. To simplify the mathematical analysis we have reduced these dynamics to a single-reset event at spike time according to eq. (8). This simplification is natural given an approximately constant duration of the active phase of an action potential, and it will become evident from our results that this simplification does not qualitatively change the dynamics of the network compared to Bose et al.’s original equations, where depression dynamics are described by an exponential decay.

When the cells are uncoupled 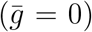, the membrane dynamics are determined by the cubic *v*-nullcline *v*_∞_(*v_i_*) and the sigmoid *w*-nullcline *w*_∞_(*v_i_*), satisfying 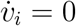 and 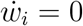, respectively. The two curves intersect along the middle branch of *v*_∞_, creating an unstable fixed point *p_f_* = (*v_f_, w_f_*) with a surrounding stable limit cycle of period *T* (fig. 1A). Trajectories along that limit cycle have the familiar shape of the action potential (fig. 1B). The trajectory of an action potential can be dissected into four phases: (1) a silent phase, (2) a jump up, (3) an active phase, and (4) a jump down (see e.g. Ermentrout & Terman, 2010). During the silent phase the trajectory evolves along the left branch (*v_i_* < *v_θ_*) of the cubic *v*-nullcline. Once the trajectory reaches the local minimum of *v*_∞_, it “jumps up” to the right branch (*v_i_* > *v_θ_*), crossing the firing threshold *v_θ_*. During the active phase the trajectory then evolves along the right branch of the cubic until it arrives at the local maximum, where it “jumps down” to the left branch commencing a new cycle.

**Figure 1:**
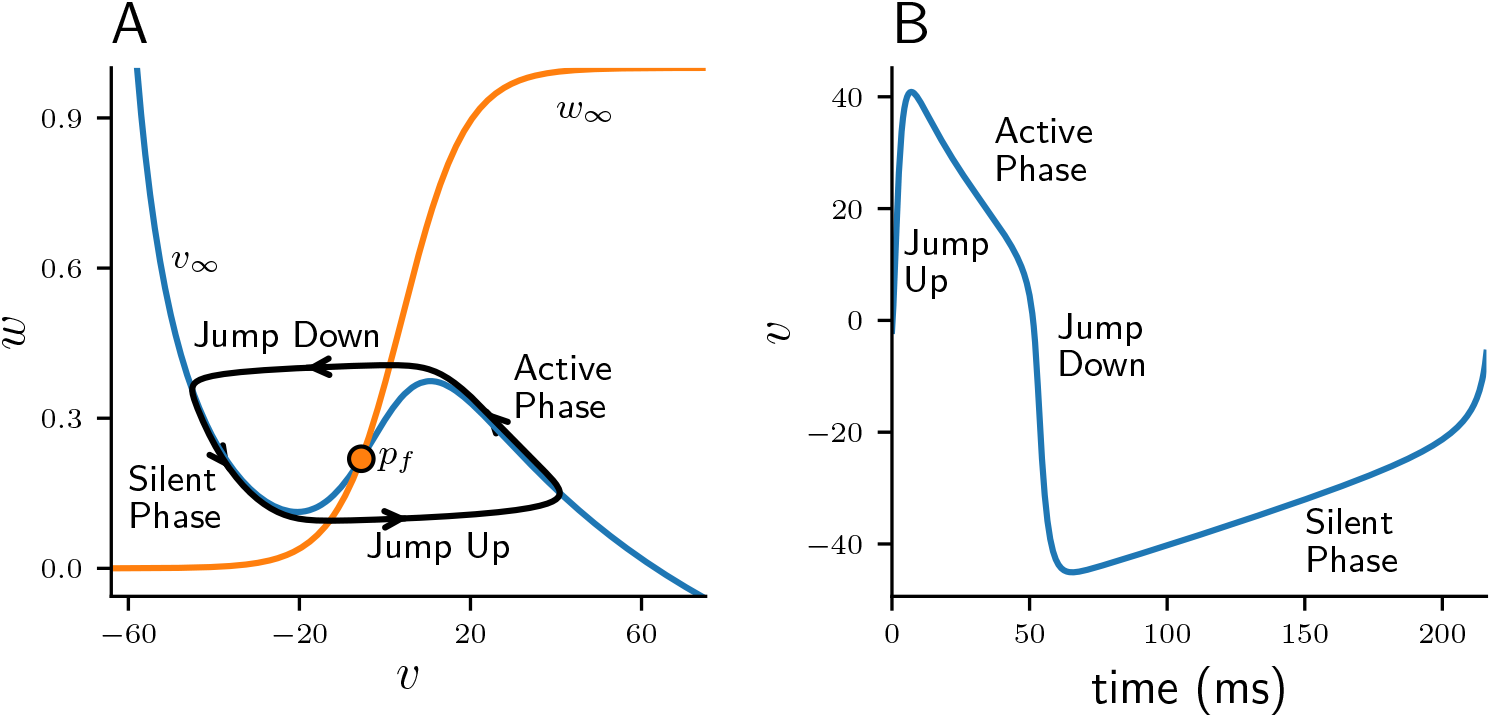
Periodic solution of relaxation-oscillator model neuron. **A**: Projection of limit cycle onto (*v, w*)-phase plane with *v*-nullcline (blue, *v*_∞_) and *w*-nullcline (orange, *w*_∞_). Unstable fixed point *p_f_* is indicated by an orange dot. **B**: Corresponding voltage trace *v*(*t*) of an action potential.

The two-cell network model is numerically integrated using an adaptive step-size integrator for stiff differential equations implemented with XPPAUT (Ermentrout, 2002) and controlled through the Python packages SciPy (Virtanen et al., 2020) and PyXPP (Olenik, 2021). The following mathematical analysis is performed on the equations of a single cell. Unless required for clarity, we will therefore omit the subscripts *i, j* from here on.

## 3. Results

### 3.1. Anti-phase burst solutions

Short-term synaptic depression of inhibition in a half-centre oscillator acts as a *burst termination* mechanism (Brown, 1911) and is known to produce *n* – *n* anti-phase burst solutions of varying period. Such *n* – *n* solutions consist of cells firing bursts of *n* spikes in alternation. Figure 2A shows a typical 4 – 4 burst. While one cell is firing a burst it provides an inhibitory conductance to the other cell, preventing it from firing. Therefore, at any given moment one cell is spiking while the other is inhibited. Consistent with Bose & Booth (2011) we will refer to the currently firing cell as “free” and we will call the inhibited cell “quiet”. Additionally, we will distinguish between two phases of a *n* – *n* solution: We will refer to the burst duration of a cell as the “free phase”, which is the time between the first spike and the last spike in a burst. And we will call the remaining duration of a cycle, when a cell is not spiking, the “quiet phase”.

**Figure 2:**
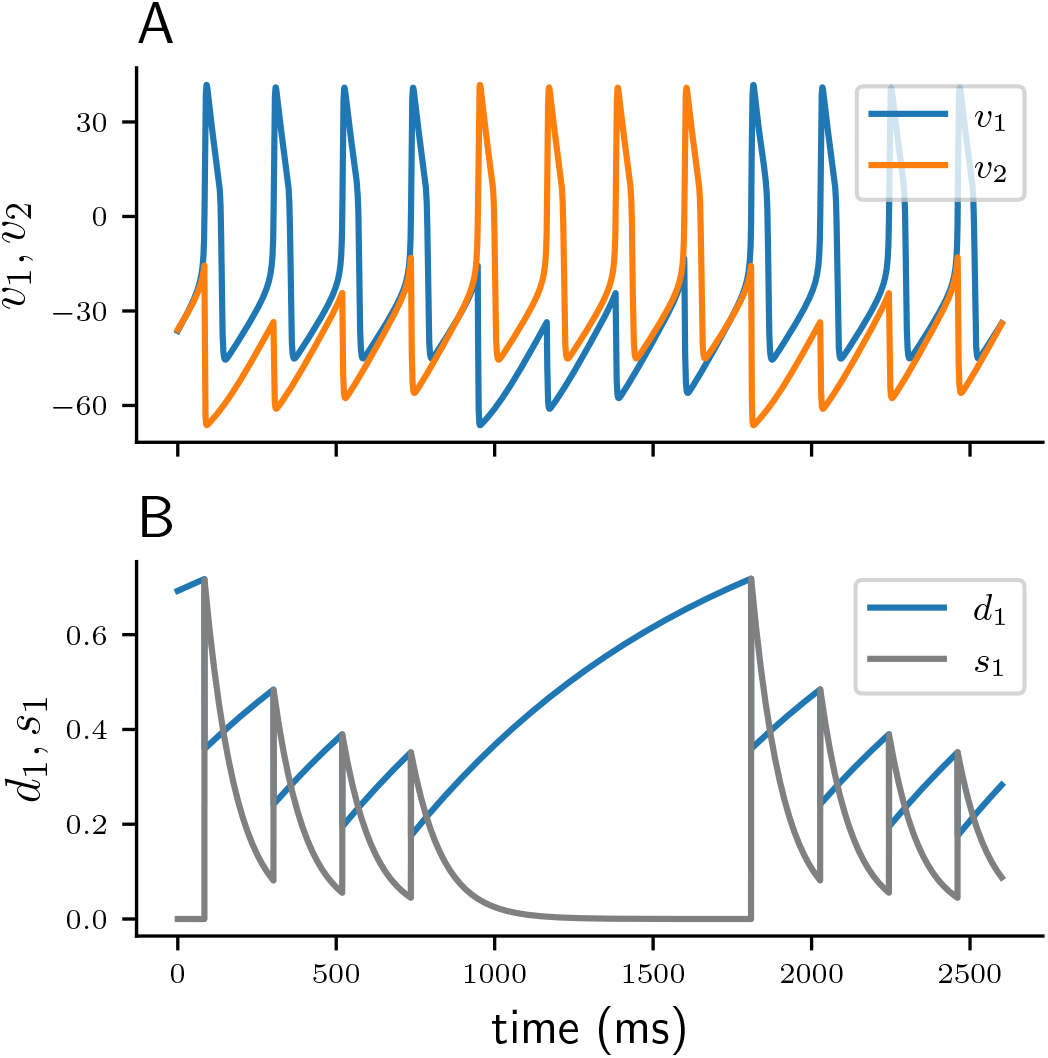
Solution profiles of a 4 – 4 burst. **A**: Membrane potentials of cell 1 (*v*_1_, blue), and cell 2 (*v*_2_, orange). **B**: Synaptic variables *d*_1_ (blue) and *s*_1_ (grey) of cell 1.

With each action potential of the free cell, short-term depression leads to a step-wise decrease of *d*, and consequently of *s* (fig. 2B). If *d* depresses faster at spike time than it can recover in the inter-spike-intervals (*ISI*s), the total synaptic conductance 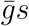 will eventually become sufficiently small to allow for the quiet cell to be released and start firing, thus inhibiting the previously free cell. While a cell is quiet its depression variable can recover. Once the quiet cell becomes free again its synaptic inhibition will be sufficient to terminate the burst of the previously free cell and commence a new cycle. As previously demonstrated by Bose & Booth (2011), in a two-cell reciprocally inhibitory network with synaptic depression the coupling strength 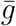 determines the type of *n* – *n* solution. Increasing 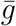 produces higher *n* – *n* burst solutions with more spikes per burst and a longer cycle period. Figure 3 shows numerically stable *n* – *n* solutions for varying values of 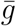. For small values of 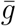 the network produces anti-phase spiking 1 – 1 solutions. As 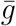 is increased the network generates solutions of increasing *n*, that is 2 – 2, 3 – 3, and 4 – 4. When 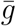 is sufficiently large (bottom of fig. 3), one of the cells continuously spikes at its uncoupled period *T* while the other cell remains fully suppressed. Depending on the initial conditions either of the two cells can become the suppressed cell, which is why the suppressed solution is numerically bistable.

**Figure 3:**
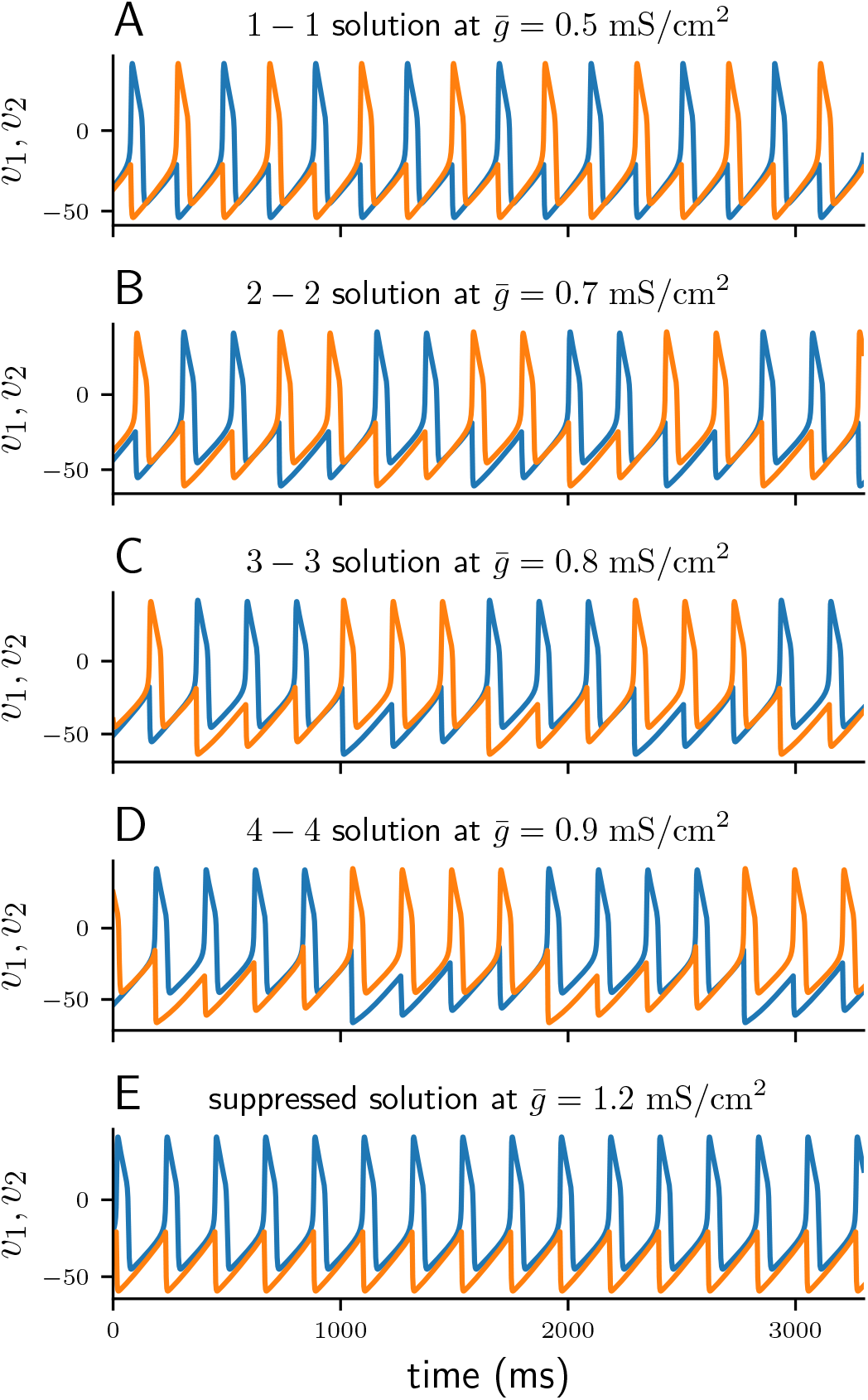
Voltage traces of numerically stable solutions for increasing values of the coupling strength 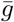 (increasing top to bottom).

Branches of numerically stable *n* – *n* solutions and their associated limit cycle period for varying values of 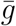 are depicted in fig. 4A (see appendix Appendix B for algorithm description). Not only do higher *n* – *n* solutions branches require stronger coupling 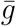, but also within *n* – *n* branches the period increases with 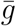. In line with Bose & Booth (2011) we find small overlaps between solution branches indicating numerical bistability, for example such as between the 2 – 2 and 3 – 3 solution branches. Branches of higher *n* – *n* burst solutions occur on increasingly smaller intervals of 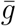, for instance is the 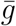 interval of the 5 – 5 branch shorter than that of the 4 – 4 branch. The interval between the 5 – 5 branch and the suppressed solution (region between dotted lines in fig. 4A) not only contains even higher numerically stable *n* – *n* solutions, such as 11 – 11 bursts, but also other non-symmetric *n* – *m* solutions as well irregular, non-periodic solutions. However, the analysis in the following sections will only be concerned with the numerically stable and symmetric *n* – *n* solutions.

**Figure 4:**
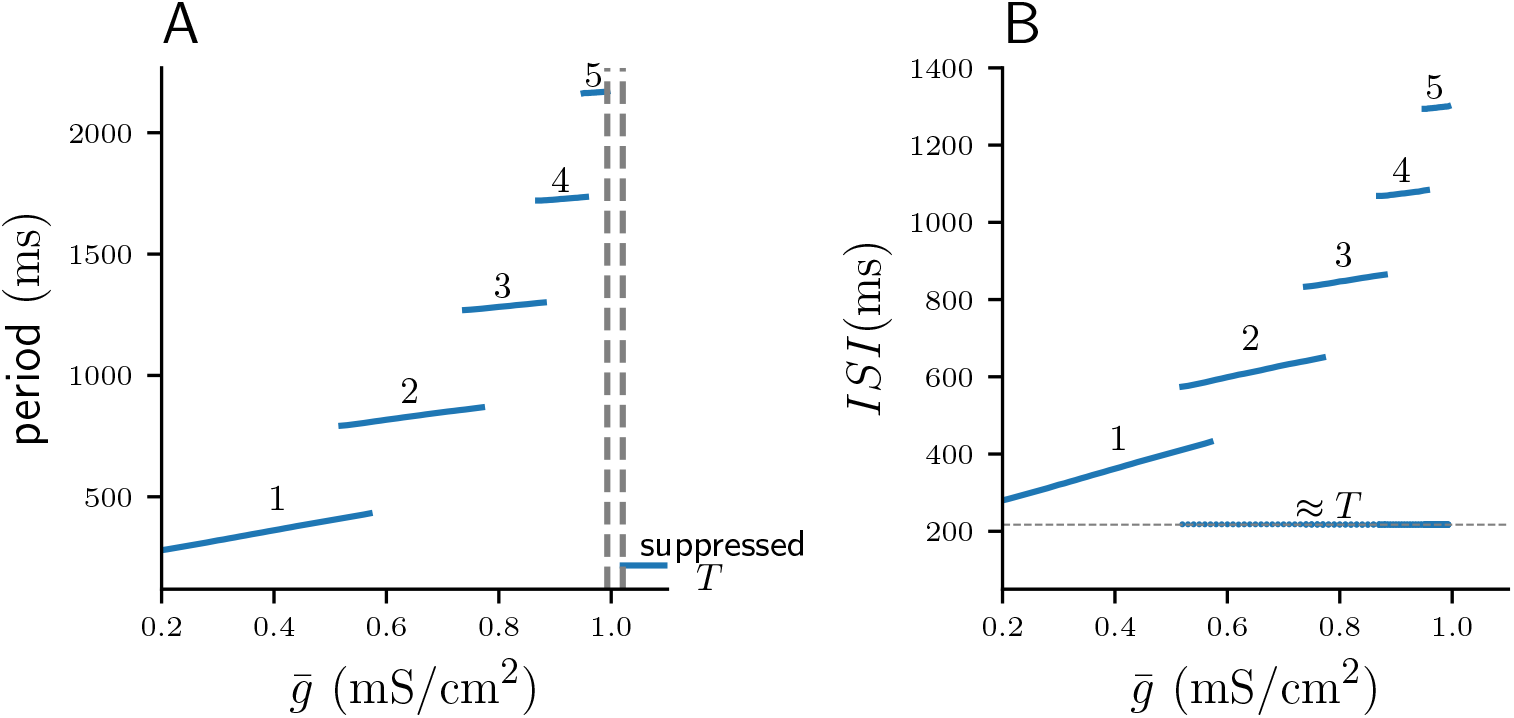
Numerically computed bifurcation diagrams of stable *n* – *n* solutions for increasing coupling strength 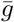. **A**: Period of stable solutions. Dashed lines show the interval between the 5 – 5 and the suppressed solutions, where higher period *n* – *n* solutions occur on increasingly smaller intervals of 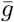. **B**: *ISI*s corresponding to *n* – *n* solutions in A. Long *ISI*s are associated with the quiet phase of a burst, short *ISI*s with the free phase. During the free phase, *ISI*s are of approximately constant duration *T*.

### 3.2. Mathematical analysis of two-cell network

The goal of the following mathematical analysis is to reduce the complexity of the eight-dimensional system to some easily tractable quantity. As we will see later this quantity is the value of the depression variable *d* of either of the two cells. We will construct the solution of *d* in a piecewise manner from one spike to the next, first during the free phase, and then during the quiet phase. This construction will require two assumptions about the membrane and synaptic dynamics. The first assumption states that during a burst the free cell fires at its uncoupled period *T*, which allows us to build the solution of *d* during the free phase. The second assumption states that once the inhibitory conductance acting on the quiet cell drops below a critical threshold, the cell is immediately released and fires. The second assumption is necessary to predict the release time of the quiet cell, which allows us to model the recovery of *d* during the quiet phase. In other words, the second assumption requires that the release of the quiet cell from inhibition depends only on the timecourse of the inhibition, and not on the membrane dynamics of the quiet cell. Both assumptions can be observed in coupled relaxation-oscillator types of neurons such as the Morris-Lecar model we use, and will be numerically verified below. Both assumptions were first explored in Bose & Booth (2011) to derive algebraic conditions that guarantee the periodicity of the depression variable for different *n* – *n* solutions. However here we will use these assumptions to construct a Poincaré map of *d*, which will provide a geometric intuition for the dynamics of the full two-cell network and its dependence on model parameters.

Our first assumption about the model states that the free cell fires at its uncoupled period *T*, that is, during the free phase of a burst we have *ISI* = *T*. Solution profiles in fig. 3 suggest that the *ISI*s in the free phase are indeed approximately constant. We can further numerically confirm this observation by capturing the *ISI*s of the stable solutions from the bifurcation diagram in fig. 4A. Figure 4B shows two types of *ISI*s as 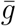 is varied, long *ISI*s and short *ISI*s. Long *ISI*s lie on multiple branches, each branch associated with a stable *n* – *n* solution, and are monotonically increasing with 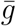. These *ISI*s lie in the quite phase of the burst, and thus correspond to the time interval between the last spike time of a burst, and the full cycle period. The short *ISI*s are calculated from the spikes within the free phase and do not vary significantly with 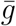, but are approximately *ISI* ≈ *T*, affirming our first assumption. Assuming *ISI* = *T* allows us to ignore the non-linear membrane dynamics during the free phase, and to construct the evolution of the synaptic variables iteratively from spike to spike. Assuming *ISI* = *T* in the free phase seems reasonable given that inhibition acting on the quiet cell decays exponentially to zero on a much shorter timescale than the duration of the *ISI*, and therefore, once the quiet cell is released its trajectory quickly approaches the spiking limit cycle.

Our second assumption states that the quiet cell is released and spikes as soon as inhibition from the free cell drops below a constant threshold. We will now define such a “release condition” by exploiting the discrepancy in timescales between the fast membrane dynamics, and the slower synaptic dynamics. Let us first consider the dynamics of a single Morris-Lecar neuron. We fix the synaptic variable that acts on the cell by setting *s* = 1, which also makes the applied synaptic conductance 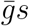 constant. Recall from fig. 1A that in case of a single uncoupled cell 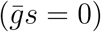, the *v*- and *w*-nullclines intersect at some unstable fixed point *p_f_* = (*v_f_*, *w_f_*), while trajectories revolve around a stable spiking limit cycle. Increasing 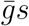 moves the cubic *v*_∞_ with the ensuing unstable fixed point *p_f_* down the sigmoid *w*_∞_ in the (*v* – *w*)-plane (fig. 5). When 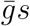 is large enough, the fixed point *p_f_* becomes stable, attracting all previously periodic trajectories. There exists a unique value 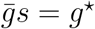 when *p_f_* changes stability and the stable limit cycle vanishes. Thus, when a constant inhibitory conductance is applied, 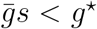 acts as a necessary condition for a cell to spike. In contrast, when 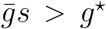 inhibition is strong enough to prevent a cell from spiking (Bose & Booth, 2011).

**Figure 5:**
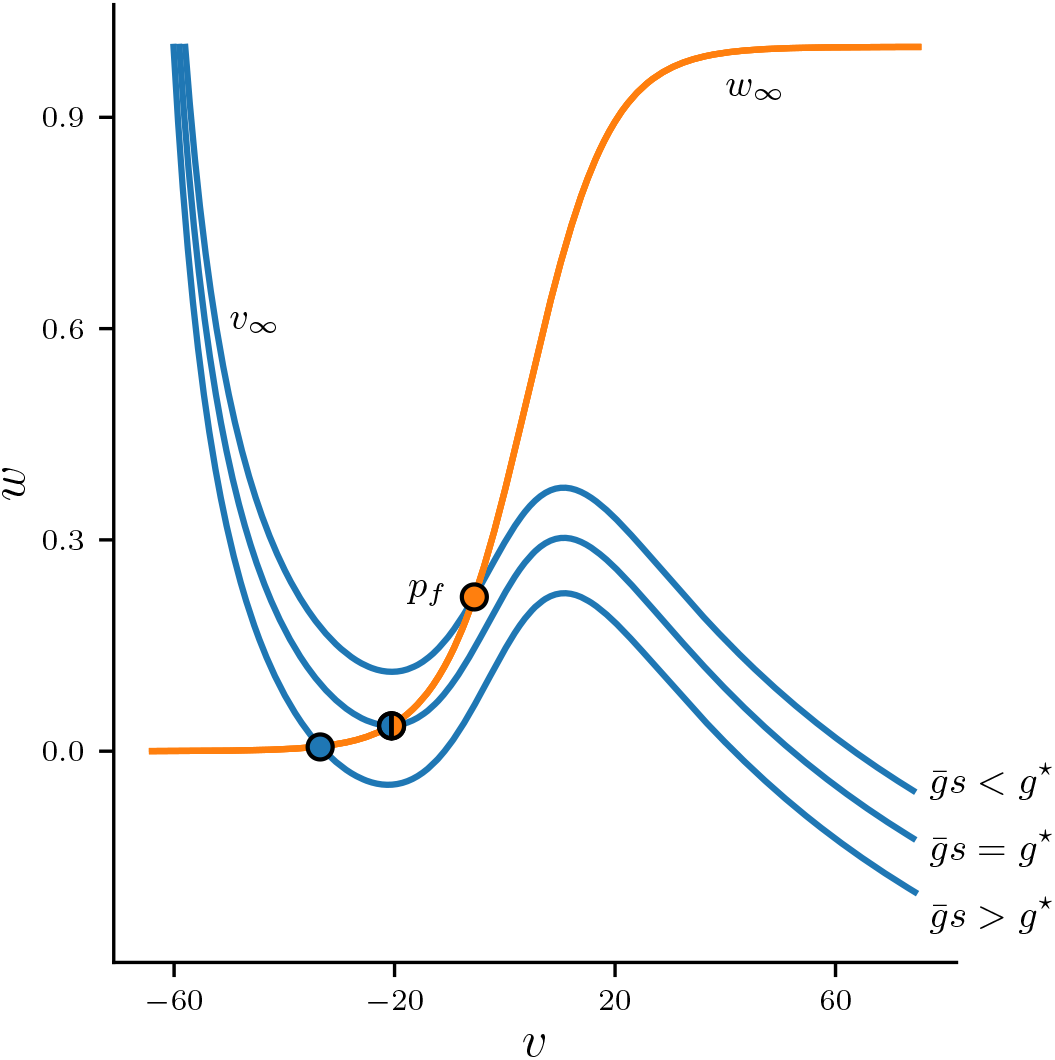
Nullclines *v*_∞_ (blue) and *w*_∞_ (orange) in the (*v*, *w*)-phase plane for different values of the total synaptic conductance 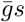. For small 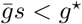, fixed point *p_f_* is unstable (orange point). Larger values 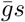 move *v*_∞_ down in the (*v*, *w*)-plane until *p_f_* changes stability (half orange, half blue point) at some critical total conductance value 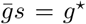, and becomes stable (blue point) for 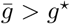.

Now let us analyse the nullclines of the quiet cell when the two cells are coupled via synaptic inhibition with depression. Let 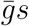 here denote the total synaptic conductance which acts on the quiet cell and is produced by the free cell, and let *p_f_* be the fixed point associated with the quiet cell. At the start of the burst of the free cell we have 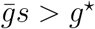 and *p_f_* is stable. When the free cell spikes 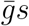 peaks (eq. (7)) and *p_f_* jumps down the *w*-nullcline. Then in between spikes 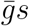 decays exponentially (eq. (5)) causing *p_f_* to move up the *w*-nullcline while attracting trajectories of the quiet cell. Once depression causes the synaptic conductance to become small enough to satisfy 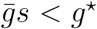 and the quiet cell is released, fixed point *p_f_* becomes unstable allowing the quiet cell to fire. If the trajectory of the quiet cell remains sufficiently close to *p_f_* when it changes stability, then 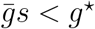 acts as a release condition that is not only necessary, but also sufficient for firing of the quiet cell. In this case the release of the quiet cell occurs precisely when

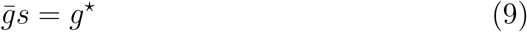

is satisfied.

Whether the (*v, w*)-trajectory of the quiet cell can remain close enough to *p_f_* to make eq. (9) sufficient for firing depends largely on the coupling strength 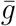 and the timescale disparity between membrane dynamics and synaptic dynamics (Bose & Booth, 2011). It is straightforward to test our assumption of a release condition by numerically integrating the full system of ODEs and calculating the time interval between the first spike of the quiet cell and the time when 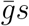 first crosses *g*^⋆^. We will call this time interval the “release delay”. If our assumption holds, we would expect an approximately zero release delay. Figure 6 shows the numerically computed graph of the release delay for varying 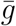. The graph shows three distinct branches, next to each branch we also plot the timecourse of a corresponding sample solution of the total synaptic conductance 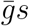 of both cells. For the rightmost branch where 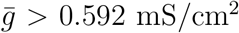 the release delay is approximately zero. Here the first spike of the quiet cell can be accurately predicted by the release condition in eq. (9). The leftmost and middle branches for 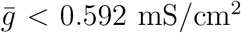 show a release delay greater than zero, the quiet cell does not immediately fire when the release condition is first satisfied, and eq. (9) does not accurately predict the release of the quiet cell. The leftmost and middle branches contain 1 – 1 and 2 – 2 solutions respectively. In both cases the coupling 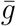 is not sufficiently strong to allow trajectories of the quiet cell to be close enough to *p_f_* to guarantee spiking once 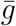 crosses *g*^⋆^. Note that in the middle branch 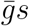 crosses *g*^⋆^ twice, and only after the second crossing does the quiet cell fire. The following map construction relies on the assumption that the release condition in eq. (9) can accurately predict the release time of the quiet cell. Given our model parameters this is only possible for sufficiently large 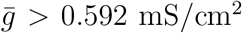. For completeness, however, we will also consider values 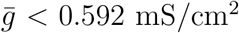 in the following analysis, bearing in mind that in this parameter range our map will not be accurate.

**Figure 6:**
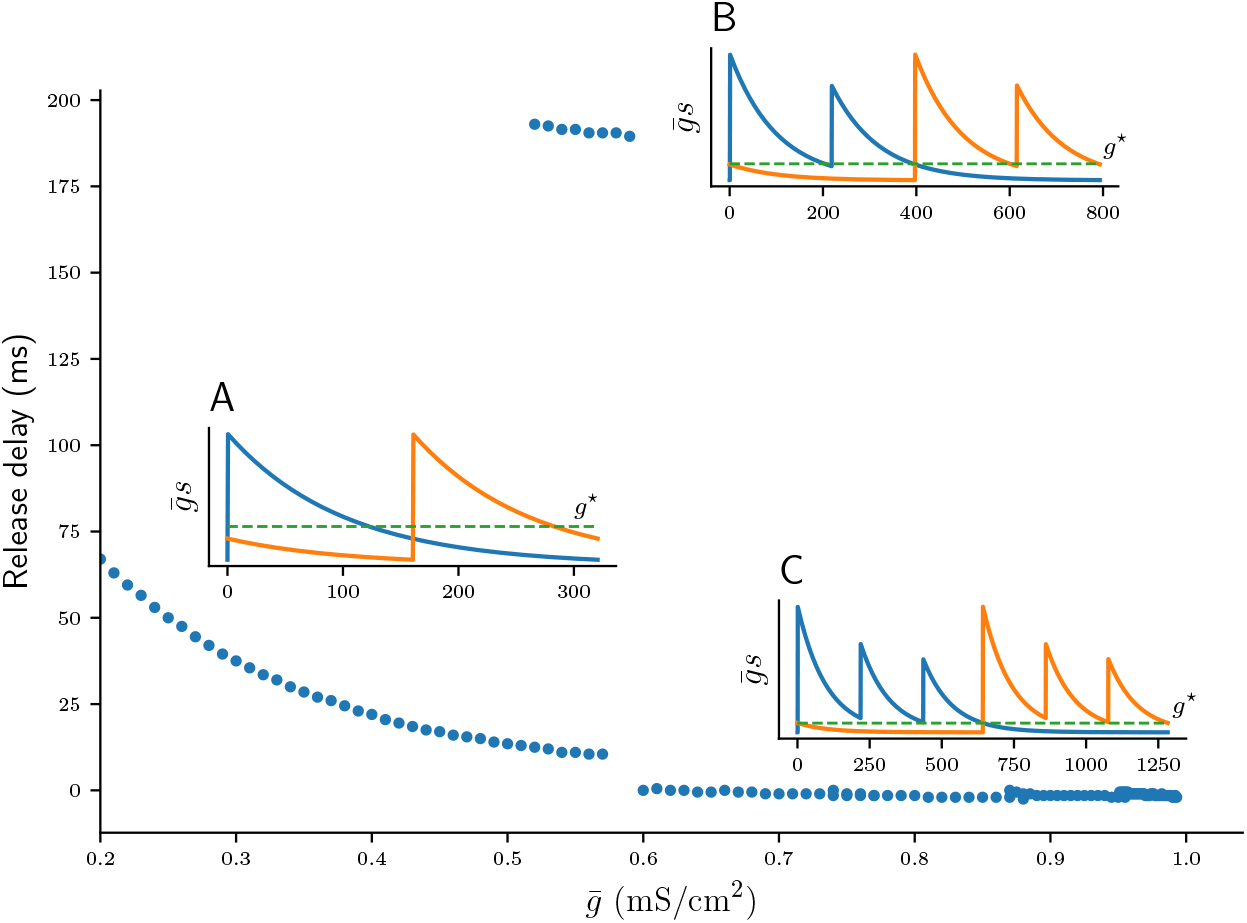
Numerically computed values of the release delay for varying 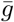. Each of the three branches (A, B, C) also shows the timecourse of the total synaptic conductance 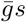 of a sample stable solution of both cells (blue and orange), as well as the release conductance *g*^⋆^ (dashed green line). **A**: Branch corresponding to the 1 – 1 solution. Here the quiet cell only spikes after a significant release delay. **B**: Branch with a long release delay associated with a subset of 2 – 2 solutions. Here the release condition is briefly satisfied after the first spike of cell 1. This does not cause firing of cell 2, which only occurs after the second spike of cell 1. **C**: Branch with *n* – *n* solutions where release delay is approximately zero and the release condition is sufficient for firing of cell 2.

In summary: For 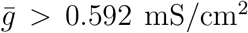 the release condition is sufficient to predict when the quiet cell is released. Due to the symmetry of *n* – *n* solutions the release occurs at exactly half the period of the full cycle, that is at *P*/2. The release time therefore uniquely determines the type of *n* – *n* solution. Furthermore, computation of the release time does not depend on the membrane nor the synaptic dynamics of the quiet cell. Instead, the solution of the synaptic variable *s* of the free cell is sufficient to predict when 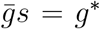 is satisfied. Finally, *s* is solely determined by the evolution of the depression variable *d* of the free cell. Constructing a solution of *d* during the free phase of either cell will therefore uniquely determine the solution of the full eight-dimensional network. However, finding the solution *d* requires us to know the initial value *d*(0) at the start of a cycle at *t* = 0. In the next section we will construct a scalar return map that tracks these initial values *d*(0) from cycle to cycle of stable *n* – *n* solutions.

### 3.3. Construction of the scalar Poincaré map ∏_n_

In this section we construct the scalar Poincaré map ∏_*n*_ : *d*^⋆^ ↦ *d*^⋆^. Here the discrete variable *d*^⋆^ tracks the values of the continuous depression variable *d* at the beginning of each *n* – *n* burst. The map ∏_*n*_ therefore describes the evolution of *d*, of either of the two cells, from the beginning of one cycle to the beginning of the next cycle. To simplify the map construction we will assume that a free cell fires exactly *n* times before it becomes quiet. Later we will relax this assumption. We will construct ∏_*n*_ by evolving *d* first during the free phase and then during the quiet phase of the *n* – *n* limit cycle. First, let us give explicit definitions of the free and quiet phases. A schematic illustration of both phases is given in fig. 7.

**Figure 7:**
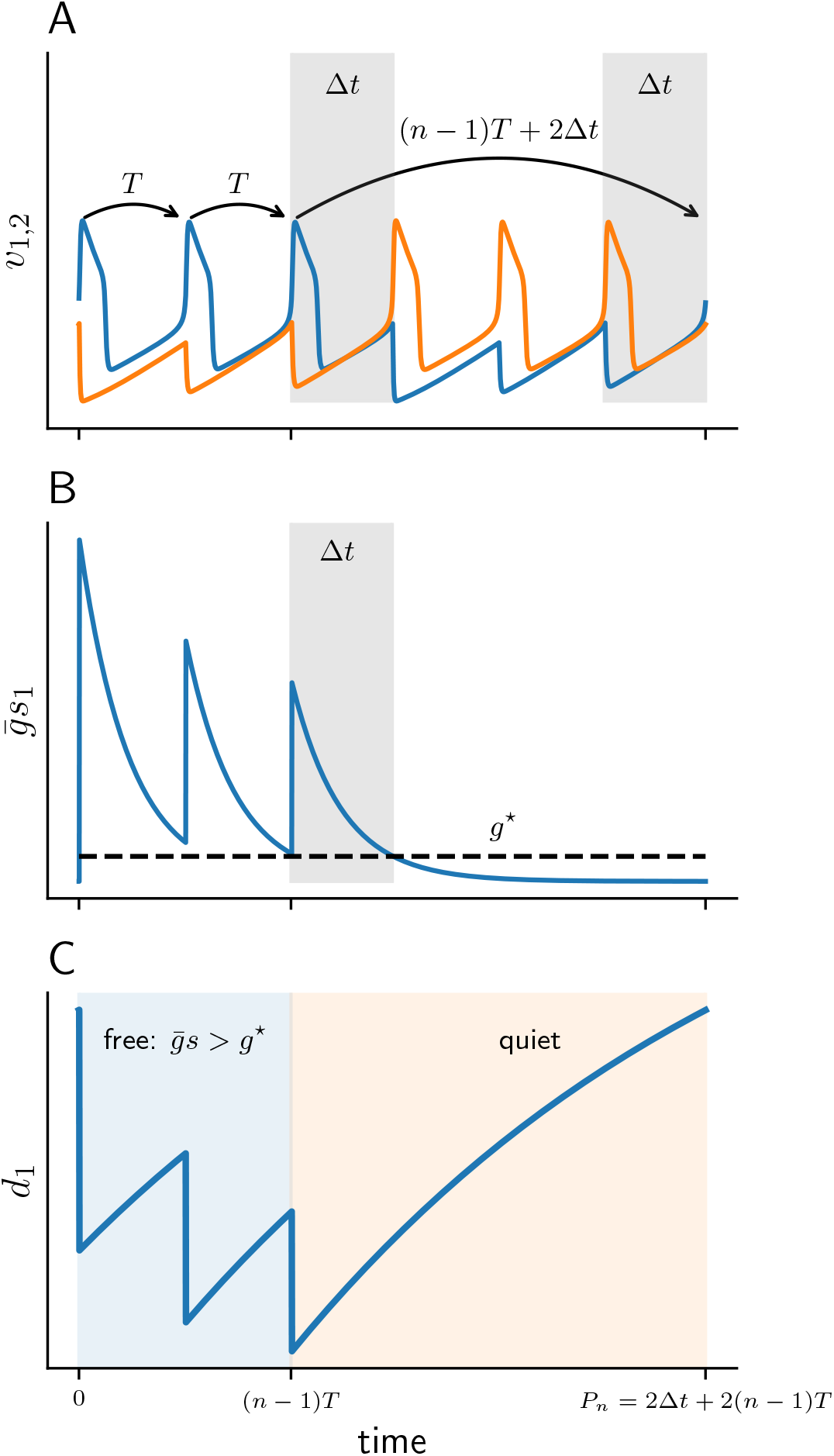
Schematic diagram of the free and quiet phases for a 3–3 solution. **A**: Membrane potentials of cell 1 (*v*_1_) and cell 2 (*v*_2_). The grey patches depict inter-burst-intervals Δ*t*. **B**: Total synaptic conductance of cell 1 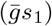 as it crosses the release conductance *g*^⋆^. **C**: Solution *d*_1_(*t*) of depression variable of cell 1, during free (blue) and quiet phases (orange).

Suppose that at *t* = 0 cell 1 becomes free with some initial *d*(0). Cell 1 then fires *n* spikes at the uncoupled period *T*. Let *s*(*t*) and *d*(*t*) be the corresponding solutions of the synaptic and depression variables of cell 1. After *n* spikes the total conductance 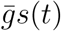 acting on the quiet cell 2 has decayed sufficiently to satisfy the release condition (9), that is at some time *t* = (*n* – 1)*T* + Δ*t* we have 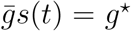. Cell 2 is then released and prevents cell 1 from further spiking. Here Δ*t* is the *inter-burst-interval*, or the time between the last spike of cell 1 and the first spike of cell 2 (Bose & Booth, 2011). Once released, cell 2 also fires *n* spikes until cell 1 becomes free once again at the cycle period. Let *P_n_* denote the full cycle period of a *n* – *n* solution:

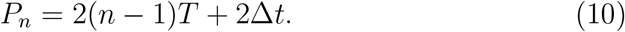

We can now define the free and quiet phases of cell 1 explicitly. The free phase is the time interval between the first and last spikes of the burst, that is for time 0 < *t* < (*n* – 1)*T*. During the free phase of cell 1, the quiet cell 2 is inhibited sufficiently strong to prevent it from firing, hence 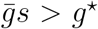. The quiet phase of cell 1 is the remaining duration of the cycle when the cell is not firing, that is for (*n* – 1)*T* < *t* < 2(*n* – 1)*T* + 2Δ*t*.

Note that only the quiet phase depends on Δ*t*, and Δ*t* will play a central role in the construction of ∏_*n*_. From eq. (10) Δ*t* can be be computed as

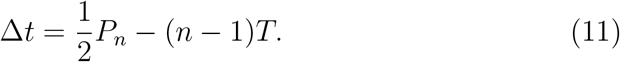

We can use eq. (11) and the numerically computed bifurcation diagram of the period for stable *n* – *n* solutions in fig. 4A to obtain the graph of Δ*t* as a function of 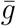 (fig. 8). Each continuous branch of Δ*t* is monotonically increasing and corresponds to a *n* – *n* burst: Stronger coupling 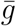 increases the total synaptic conductance 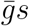 that acts on the quiet cell, thus delaying its release. It is easy to see that for any *n*-branch we have Δ*t* < *T*: Once Δ*t* crosses *T*, the free cell can “squeeze in” an additional spike and the solutions bifurcate into a (*n* + 1) – (*n* + 1) burst.

**Figure 8:**
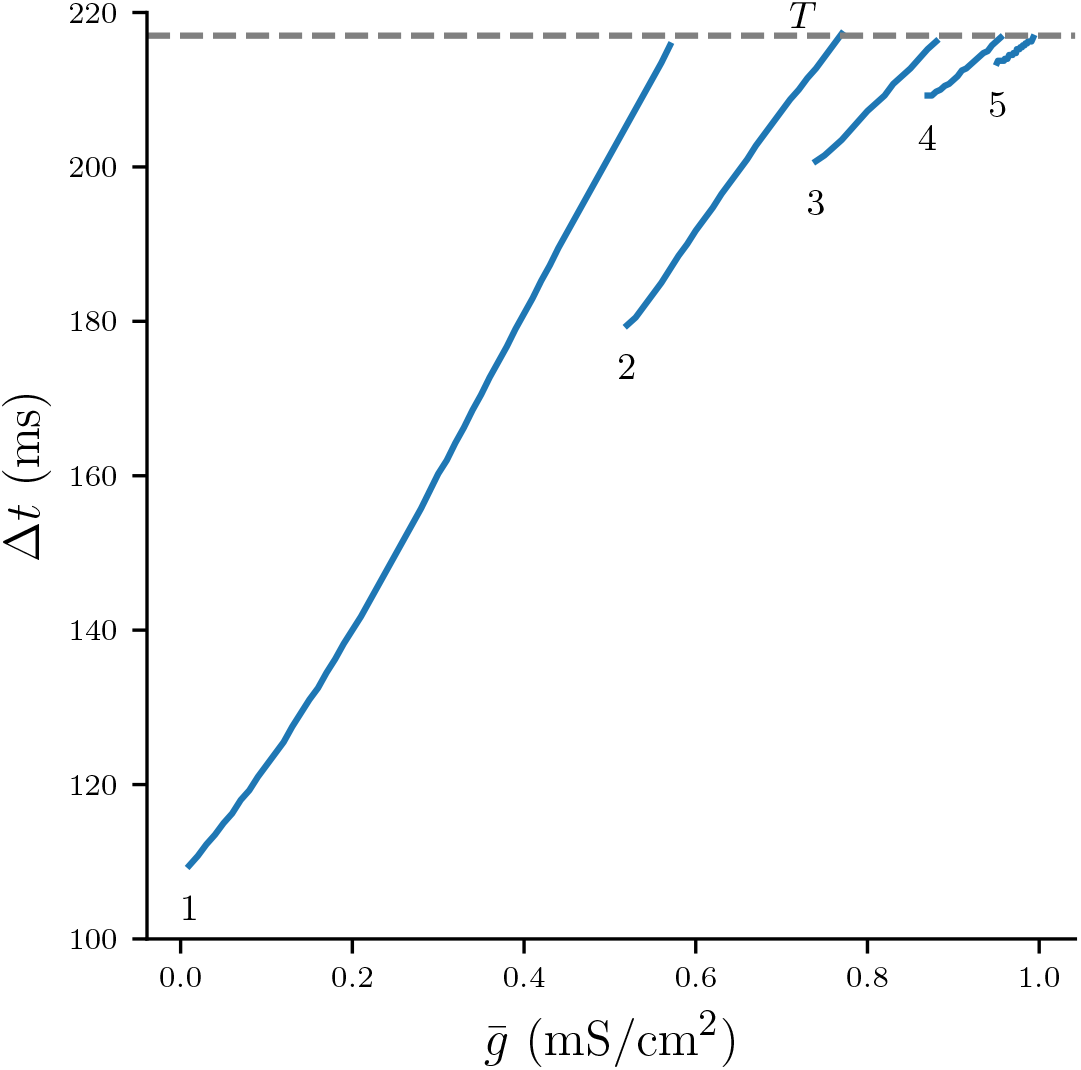
Numerically computed bifurcation diagram of Δ*t* for varying 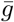. Each continuous branch is associated with a stable *n* – *n* burst solution. Increasing 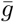 increases Δ*t* until the solutions bifurcate at Δ*t* ≈ *T*.

Distinguishing between the free and quiet phases of a cycle allows us to describe the dynamics of the depression variable *d* explicitly for each phase. As can be seen from fig. 7C, during the free phase *d* depresses at spike times and recovers in the *ISI*s. In contrast, during the quiet phase *d* only recovers and does not depress. Given the initial *d*^⋆^ = *d*(0) at the beginning of the cycle and the number of spikes in the free phase *n*, we can now construct the burst map ∏_*n*_. The map

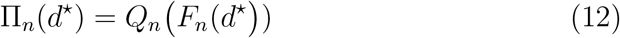

is a composition of two maps. Map

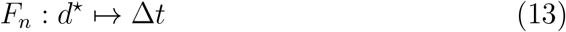

models the evolution of *d* in the free phase. *F_n_* takes an initial value *d*^⋆^ and calculates the inter-burst-interval Δ*t*. Map

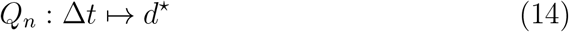

models the recovery of *d* in the quiet phase. Given some Δ*t* map *Q_n_* computes *d*^⋆^ at the start of the next cycle.

Our aim in the following analysis is to elucidate the properties of ∏_*n*_ and to understand the structure of its parameter space by exploring how the stable and unstable fixed points of ∏_*n*_ are created. To that effect it will be useful to include not only positive, but also negative values of *d*^⋆^ to the domain of ∏_*n*_. But it is important to add that values *d*^⋆^ < 0 are biologically impossible as the depression variable models a finite pool of neurotransmitters, and therefore must be positive. Because ∏_*n*_ maps first from *d*^⋆^ to Δ*t*, and then back to *d*^⋆^, we will also consider negative values of Δ*t*, interpreting them as *n* – *n* solutions with partially overlapping bursts. As will become evident, Δ*t* < 0 is only a formal violation of the biological realism of the map ∏_*n*_, as numerically stable *n* – *n* solutions of the full system of ODEs only exist for Δ*t* > 0.

We start the construction of ∏_*n*_ by first considering the free phase and building the map *F_n_*. At each spike time *t_k_*, variable *d* depresses with the reset 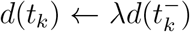 (eq. (8)). For the duration of the subsequent *ISI*, the recovery of *d* is described by the solution to eq. (6):

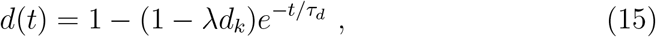

where the variable 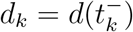 is the level of depression at spike time *t_k_*. By substituting *t* = *T* we can build a linear map that models the depression of *d* from spike time *t_k_* to the subsequent spike time *t*_*k*+1_ during the free phase:

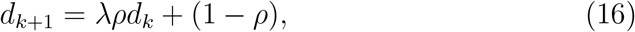

where to keep the notation simple we let *ρ* = exp(−*T*/*τ_d_*). Since 0 < λ, *ρ* < 1, the map (16) is increasing and contracting, with a fixed point at

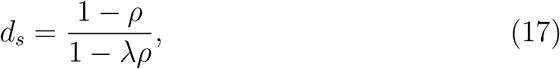

where 0 < *d_s_* < 1. The value *d_s_* is the maximum depression value that can be observed in the suppressed solution where the active cell fires at its uncoupled period *T* (see fig. 3E). Using the release condition in eq. (9) allows us to derive the value of the the minimum coupling strength that will produce the full suppressed solution, denoted as 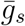. Solving eq. (5) for *s*(*t*) with *t* = *T* and setting the initial value *s*(0) = *d_s_* gives

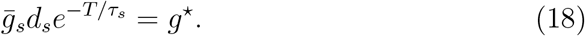

By further substituting the definition of *d_s_* in (17) and rearranging, we can write 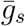 as a function of the depression strength λ:

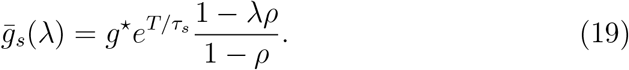

Note that the above dependence of 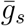 on λ is linear and monotonically decreasing. Increasing λ reduces the strength of the depression of the free cell. This in turn allows the free cell to fully suppress the quiet cell at smaller values of 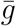.

Solving (16) gives us the linear map *δ_n_* : *d*^⋆^ ↦ *d_n_*, that for some initial *d*^⋆^ computes the depression at the *n*th spike time, 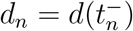:

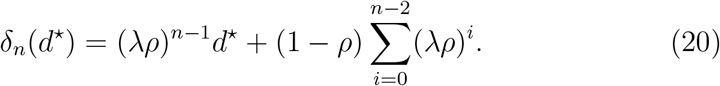

Since λ < 1, function *δ_n_* is a linearly increasing function of *d*^⋆^ with a fixed point at *d_s_* for all *n*. Having identified *d* after *n* spikes, we can now use the release condition 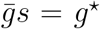 (eq. (9)) to find Δ*t*. At the last spike of the free phase at time *t_n_* = (*n* – 1)*T* the synapse variable *s* is reset according to eq. (7):

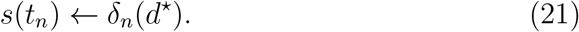

After the last spike *s* decays exponentially according to the solution of eq. (5), where setting the initial condition *s*(0) = *δ_n_*(*d*^⋆^) yields:

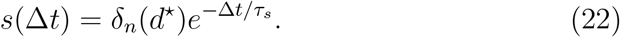

Substituting *s*(Δ*t*) into *s* of the release condition (eq. (9)) gives then

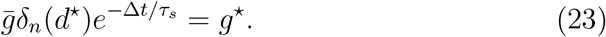

Our assumption of the release condition guarantees that the quiet cell 2 spikes and becomes free when 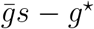 crosses zero. Solving eq. (23) for Δ*t* allows us to compute the inter-spike-interval as a function of *d*^⋆^, which defines our map *F_n_*:

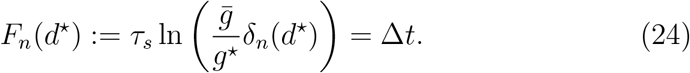

Figure 9A shows *F_n_* for various *n*, which is a strict monotonically increasing function of *d*^⋆^ as well as 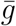. Larger values of *d*^⋆^ and 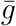, respectively, cause stronger inhibition of the quiet cell, and therefore prolong its release time and the associated Δ*t*. Map *F_n_* is defined on *d*^⋆^ > *d_a_*, where *d_a_* is a vertical asymptote found by solving *δ_n_*(*d*^⋆^) = 0 in eq. (20) for *d*^⋆^, which yields

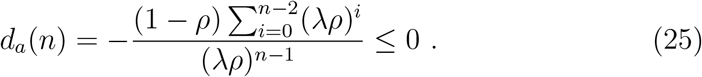

**Figure 9:**
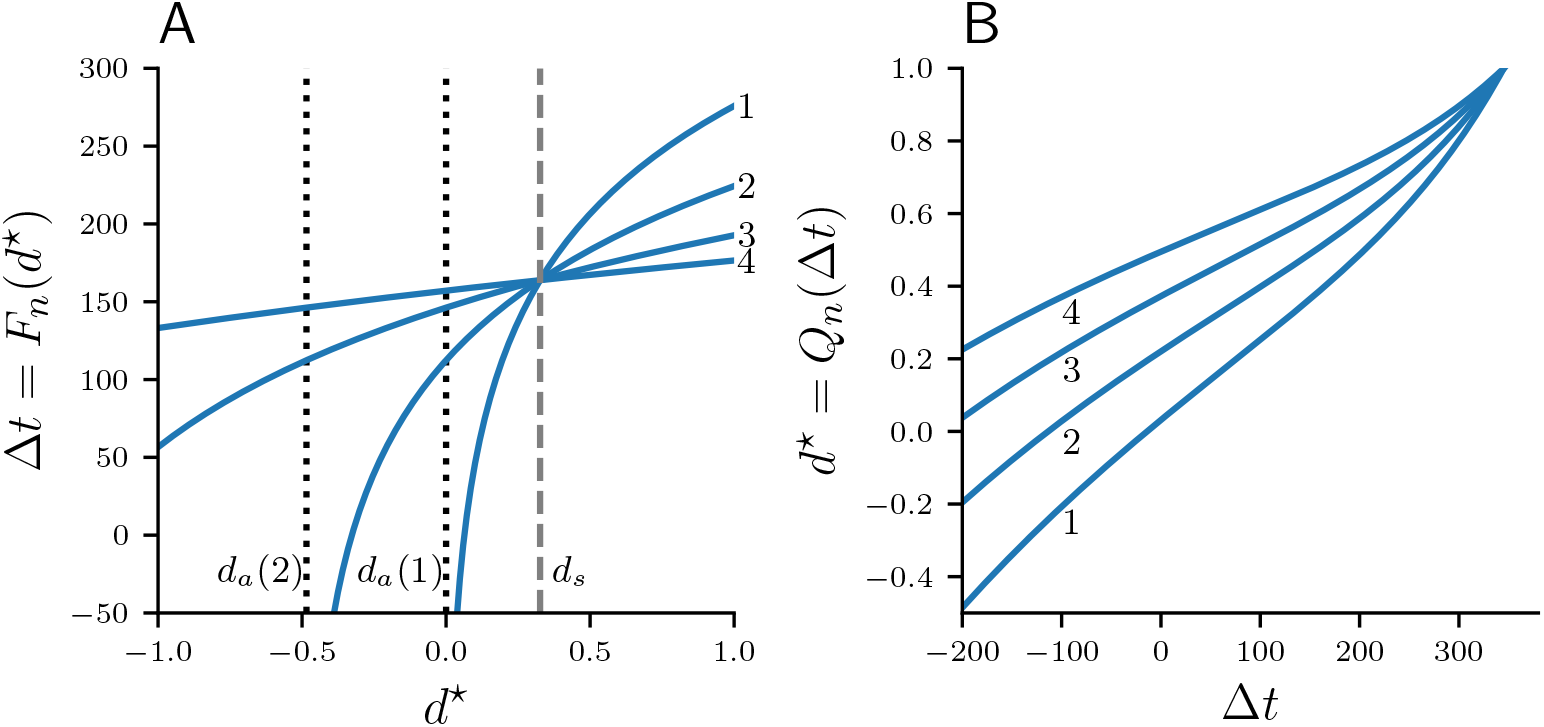
Maps *F_n_* (**A**) and *Q_n_* (**B**) for 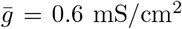 and *n* = 1, 2, 3, 4. Curves *F_n_* intersect at *d_s_* which is indicated by a dashed vertical line.

We now turn to the construction of map *Q_n_*, which describes the recovery of the depression variable during the quiet phase. As we have identified earlier, the recovery in the quiet phase of a *n* – *n* solution is of duration 2Δ*t* + (*n* – 1)*T*. Substituting that into the solution for *d*(*t*) (eq. (15)) with the initial condition *d*(0) = *δ_n_*(*d*^⋆^) yields the map *Q_n_*:

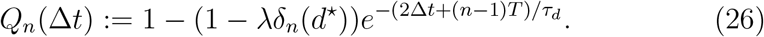

Given Δ*t*, we can find *δ_n_*(*d*^⋆^) by rearranging the release condition in eq. (23):

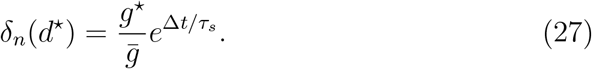

Map *Q_n_* is shown in fig. 9B for various values *n*. Note that *Q_n_* is monotonically increasing as larger values Δ*t* imply a longer recovery time, and hence *Q_n_* grows without bound. All curves *Q_n_* intersect at some 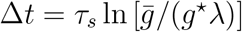 where

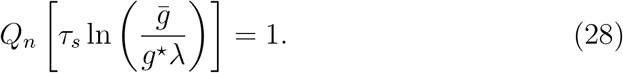

As we will show in the next section, all fixed points of the full map ∏_*n*_ occur for *d*^⋆^ < 1. We will therefore restrict the domain of *Q_n_* to 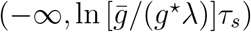 and the codomain to (−∞, 1). Additionally, while values Δ*t* > *T* will be helpful in exploring the geometry of ∏_*n*_, recall from fig. 8 that in the flow system all *n* – *n* solutions bifurcate into (*n* + 1) – (*n* +1) solutions exactly when Δ*t* = *T*, and we will address this concern in the last part of our map analysis.

Having found *F_n_* and *Q_n_*, we can now construct the full map ∏_*n*_(*d*^⋆^) = *Q_n_*(*F_n_*(*d*^⋆^)):

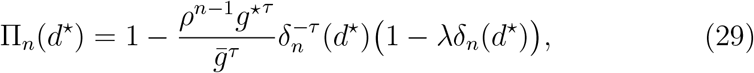

where we substituted *τ* = 2*τ_s_*/*τ_d_*. Since *d* is the slowest variable of the system and *τ_d_* ≫ *τ_s_*, we will also assume *τ* < 1. Figure 10A depicts ∏_*n*_ for various *n*. Intersections of ∏_*n*_ with the diagonal are fixed points of the map. Figure 10B shows ∏_2_ with *n* = 2. Varying the synaptic strength 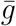 moves the curves ∏_*n*_ up and down the (*d*^⋆^, ∏_*n*_)-plane. For 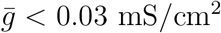 map ∏_2_ has no fixed points. As 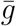 is increased to 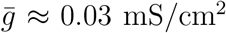, curve ∏_2_ coalesces with the diagonal tangentially. When 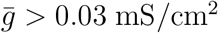, a pair of fixed points emerge, one stable and one unstable fixed point, indicating the occurrence of a fold bifurcation of maps.

**Figure 10:**
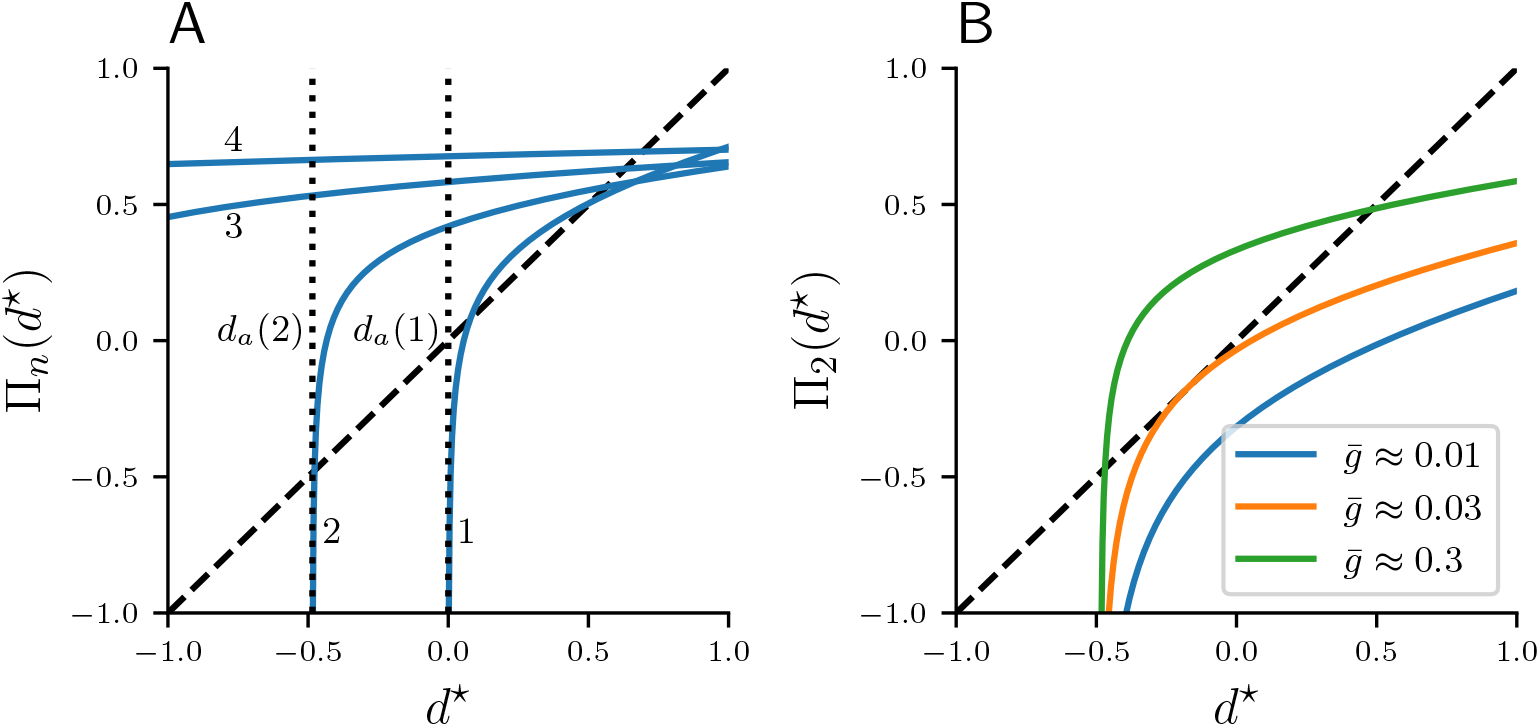
Map ∏_*n*_ : *d*^⋆^. **A**: ∏_*n*_ for *n* = 1, 2, 3, 4 at 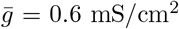. **B**: ∏_2_ with *n* = 2 for varying 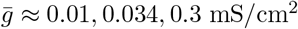. The identity function is illustrated by a diagonal line.

From eq. (29) it is evident that ∏_*n*_ is monotonically increasing with respect to 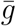 and also *d*^⋆^:

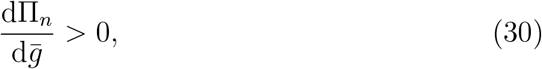

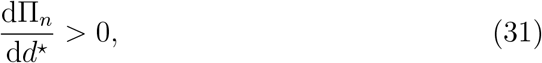

and in the following sections we will heavily rely on this monotonicity property of ∏_*n*_. Just as *F_n_*, curves ∏_*n*_ spawn at the asymptote *d_a_* (eq. (25)), and because

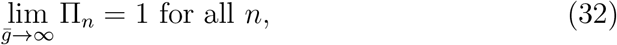

fixed points of ∏_*n*_ lie in (*d_a_*, 1).

### 3.4. Existence and stability of fixed points of ∏_*n*_

We introduce the fixed point notation 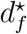 with 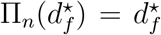. The existence of fixed points 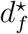 for 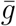 sufficiently large can be shown from the strict monotonicity of ∏_*n*_ with respect to 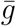 and *d*^⋆^ (eqs. (30) and (31)), as well as the fact that the slope of ∏_*n*_ is monotonically decreasing,

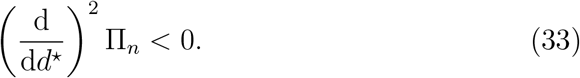

In the limit *d*^⋆^ ↦ *d_a_* the value of ∏_*n*_ decreases without bound for any 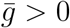. For *d*^⋆^ > *d_a_* the value of ∏_*n*_ depends on 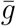. In the limit 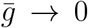, ∏_*n*_ also decreases without bound, but as 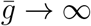 values of ∏_*n*_ approach 1. It follows from eq. (30) and the intermediate value theorem that for some 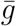 large enough ∏_*n*_ intersects the diagonal. Moreover, because ∏_*n*_ and its slope are monotonic with respect to *d*^⋆^, there exists some critical fixed point 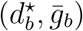 where ∏_*n*_ aligns with the diagonal tangentially with

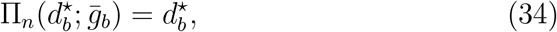

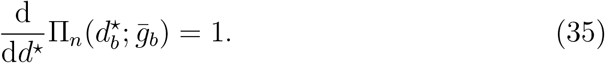

Equations (30) and (33) constitute the non-degeneracy conditions for a codimension-1 fold bifurcation of maps, indicating that in a neighbourhood of 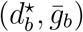 map ∏_*n*_ has the topological normal form described by the graph of

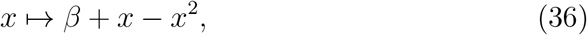

with a stable and unstable fixed point 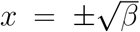, and slopes 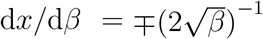, respectively.

### 3.5. Fold bifurcations of ∏_n_

Fixed points of ∏_*n*_ satisfy the fixed point equation 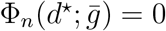, where

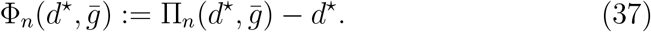

As we have already shown, for 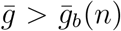 solutions to eq. (37) exist in pairs of stable and unstable fixed points. Solving eq. (37) explicitly for *d*^⋆^ it not trivial, but solving for 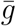 is straightforward and given by 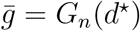, where

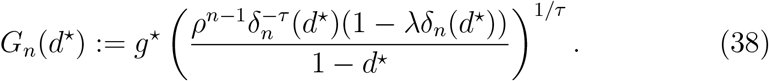

Plotting *d*^⋆^ against 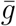 gives the fixed point curves, which are shown in fig. 11A. Note the typical quadratic shape of a fold bifurcation of maps. It is also evident that the fold bifurcations occur for increasingly smaller 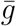 as *n* is increased. Moreover, we can observe that unstable fixed points have negative values of *d*^⋆^ for *n* > 1.

**Figure 11:**
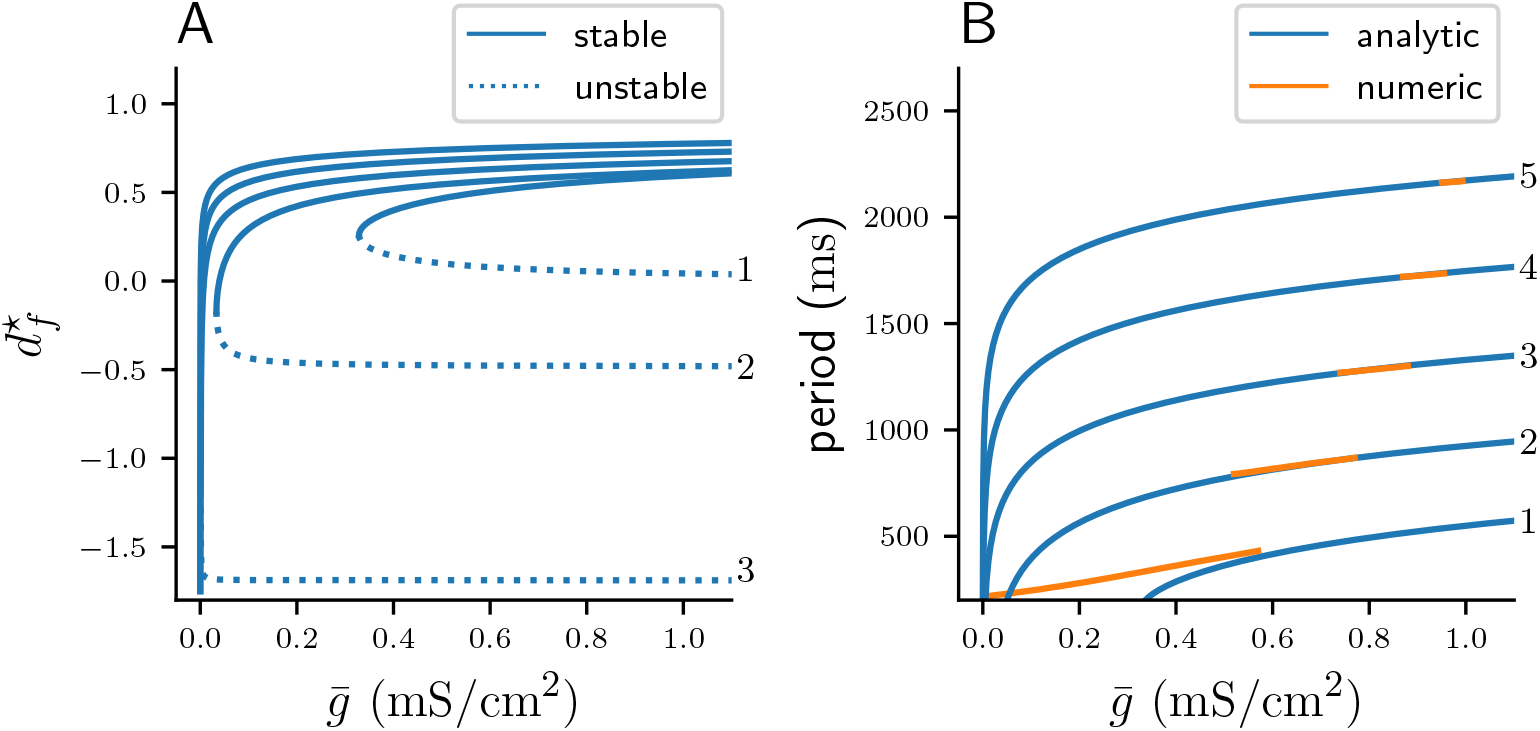
**A**: Fold bifurcation diagrams of stable (continuous curves) and unstable (dotted curves) fixed points of ∏_*n*_ for varying *n*. **B**: Cycle periods computed from stable fixed points (blue), and the corresponding solution period from numerical integration of the system of ODEs (orange).

Equation (38) also allows us to find the critical fixed point connected with the fold bifurcation, namely 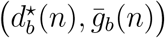, which is the global minimum of 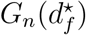:

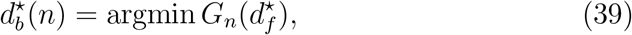

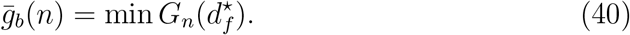

Function *G_n_* is strictly monotonic on the respective intervals of 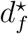 that correspond to the stable and unstable fixed points, that is

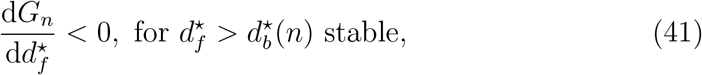

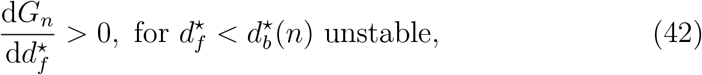

which allows us to express the stable and unstable fixed points as the inverse of *G_n_* on their respective intervals of 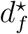. Because we are primarily interested in the stable fixed points, we define the stable fixed point function 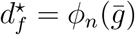 as

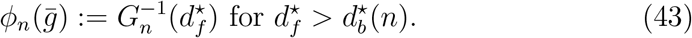

Function 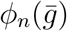 is also monotonic, and is therefore straightforward to compute numerically via root-finding. Here we use the Python package Pynverse (Gonzalez, 2021) for that purpose.

Having found the stable fixed points 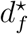 as a function of the coupling strength 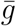, we can now compute the associated cycle period. Recall that the period is given by eq. (10), which we can be written as a function of 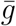:

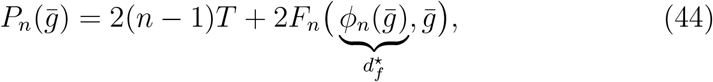

where map *F_n_* (eq. (24)) calculates the inter-burst-interval Δ*t* given a stable fixed point 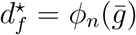. We plot the predicted period 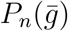 versus the cycle period that was computed from numerically integrating the full system of ODEs in fig. 11B. For *n* > 1 our map ∏_*n*_ accurately predicts the period. When laying out our assumptions in section 3.2, we have already predicted an inaccuracy for *n* =1 (see fig. 6), since here 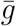 is not sufficiently strong to guarantee the validity of our release condition (eq. (9)).

It is evident from fig. 11A that *ϕ_n_* is strictly increasing with 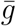. This property follows directly from the normal form of the fold bifurcation (eq. (36)), but can also be shown using implicit differentiation and the fixed point equation 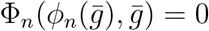 in eq. (37). For 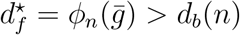 we get:

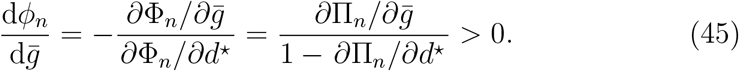

The inequality follows from 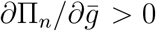 and the fact that ∂∏_*n*_/∂*d*^⋆^ < 1 for *d*^⋆^ > *d_b_*(*n*). Equation (45) allows us to explain why the period *P_n_* increases with 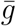, as seen in fig. 11B. Differentiating *P_n_* gives:

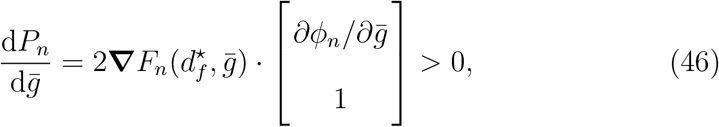

where the partial derivatives of 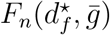 are:

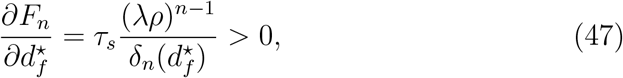

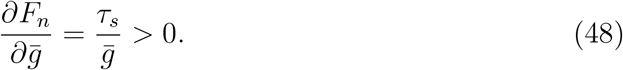

Equations (45) and (46) have an intuitive biological interpretation: Increasing the coupling strength between the neurons leads to overall stronger inhibition of the quiet cell, which delays its release and leads to a longer cycle period. The latter allows more time for the synapse to depress in the free phase and recover in the quiet phase, resulting in overall larger values of 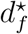, that is weaker depression at the burst onset.

While fixed points of our Poincaré map predict the cycle period of the flow system excellently, its construction relies on the strong assumption that the free phase contains exactly *n* spikes. As is evident from fig. 11B this assumption is clearly violated in the flow system, as stable *n* – *n* bursts exists only on certain parameter intervals of 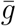. In the last sub-section we will analyse the mechanisms that guide how the stable *n* – *n* are created and destroyed, and use our previous analysis to derive the corresponding parameter intervals of 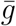 where such solutions exist.

### 3.6. Period increment bifurcations with co-existent attractors

Let 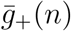 and 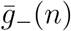 denote the left and right parameter borders on 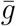 where stable *n* – *n* solutions exist. That is, as 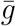 is increased stable *n* – *n* solutions are created at 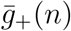 and destroyed at 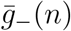. When 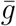 is reduced beyond 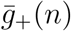, *n* – *n* solutions bifurcate into (*n* – 1) – (*n* – 1) solutions, while when 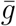 is increased beyond 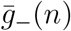, *n*–*n* solutions bifurcate into (*n*+1) –(*n*+1) solutions. Let us briefly recap our observations regarding 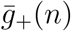 and 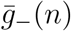 from the numerical bifurcation diagram in fig. 11B. For *n* > 1 there are the following relations:

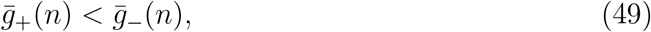

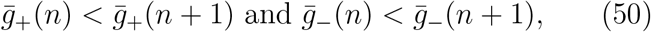

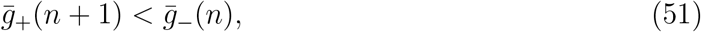

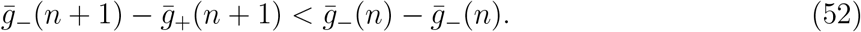

Equations (49) and (50) are self-explanatory. Equation (51) formally describes occurrence of co-existence between stable *n* – *n* and (*n* +1) – (*n* +1) solutions. Equation (52) implies that the parameter interval on 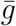 of *n* – *n* solutions decreases with *n*, in other words, bursts with more spikes occur on increasingly smaller intervals of the coupling strength. All of the above relations are reminiscent of the period-adding bifurcations with co-existent attractors, first described for piecewise-linear scalar maps with a single discontinuity by Avrutin and colleagues (e.g. see Gardini et al., 2012; Tramontana et al., 2012; Avrutin et al., 2011, 2012).

Let us now find algebraic equations that will allow us to calculate the critical parameters 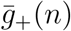 and 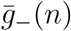 associated with the period-increment bifurcations. Recall that the period *P_n_* derived from the fixed points of ∏_*n*_ is an increasing function of 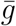:

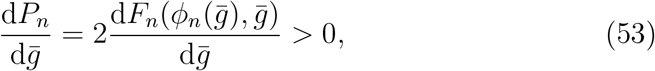

that is, as the coupling strength increases, it takes longer for the total synaptic conductance to fall below the value of the release conductance, which delays the release of the quiet cell, and Δ*t* becomes larger. Once Δ*t* = *T*, the free cell can produce another spike and the solution bifurcates into a (*n* + 1) – (*n* + 1) solution. Note, however, that at 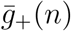 the bifurcation into a (*n* – 1) – (*n* – 1) does not occur when Δ*t* = 0. Here the mechanism is different: A sufficient reduction of 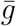 causes the total synaptic conductance to drop below the release conductance in the *previous ISI*, which allows the quiet cell to be released one spike earlier.

Using the above reasoning we can now formulate the conditions for both bifurcations at 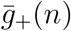 and 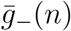. As in the previous sections, we will only restrict ourselves to the analysis of the stable fixed points given implicitly by 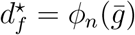 (eq. (43)). At the right bifurcation border 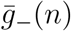 we have Δ*t* = *T*, and after substituting our *F_n_*-map (eq. (24)) this translates into

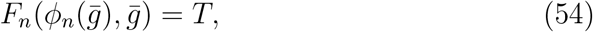

which lets us define a function

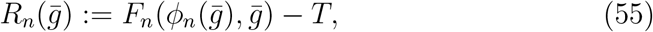

whose root is the desired right bifurcation border 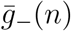. In case of the left bifurcation border at 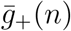 the release condition is satisfied just before the free cell has produced its *n*th spike, and after the depression variable has been reset *n* – 1 times, which gives the condition

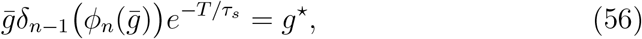

and can be rewritten as a function

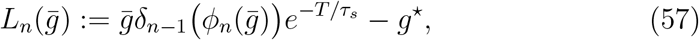

whose root is 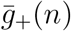. Both *R_n_* and *L_n_* are increasing with respect to 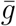, which makes finding their roots numerically straightforward.

Figure fig. 12 shows the period 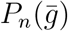 as predicted by the fixed points of ∏_*n*_ (eq. (44)) plotted on their respective intervals 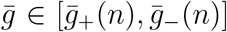 (blue), as well as the cycle period acquired from numerical integration of the full system of ODEs (orange). Note that the width of *n* – *n* branches decreases with *n*, which confirms the inequality in eq. (52). That is, bursts with more spikes occur on increasingly smaller intervals of 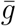, which can be interpreted as a lost of robustness with respect to the coupling strength of long-cyclic solutions. We also note the occurrence of bi-stability between pairs of *n* – *n* and (*n* + 1) – (*n* + 1) branches, which also confirms our initial observation in eq. (51).

**Figure 12:**
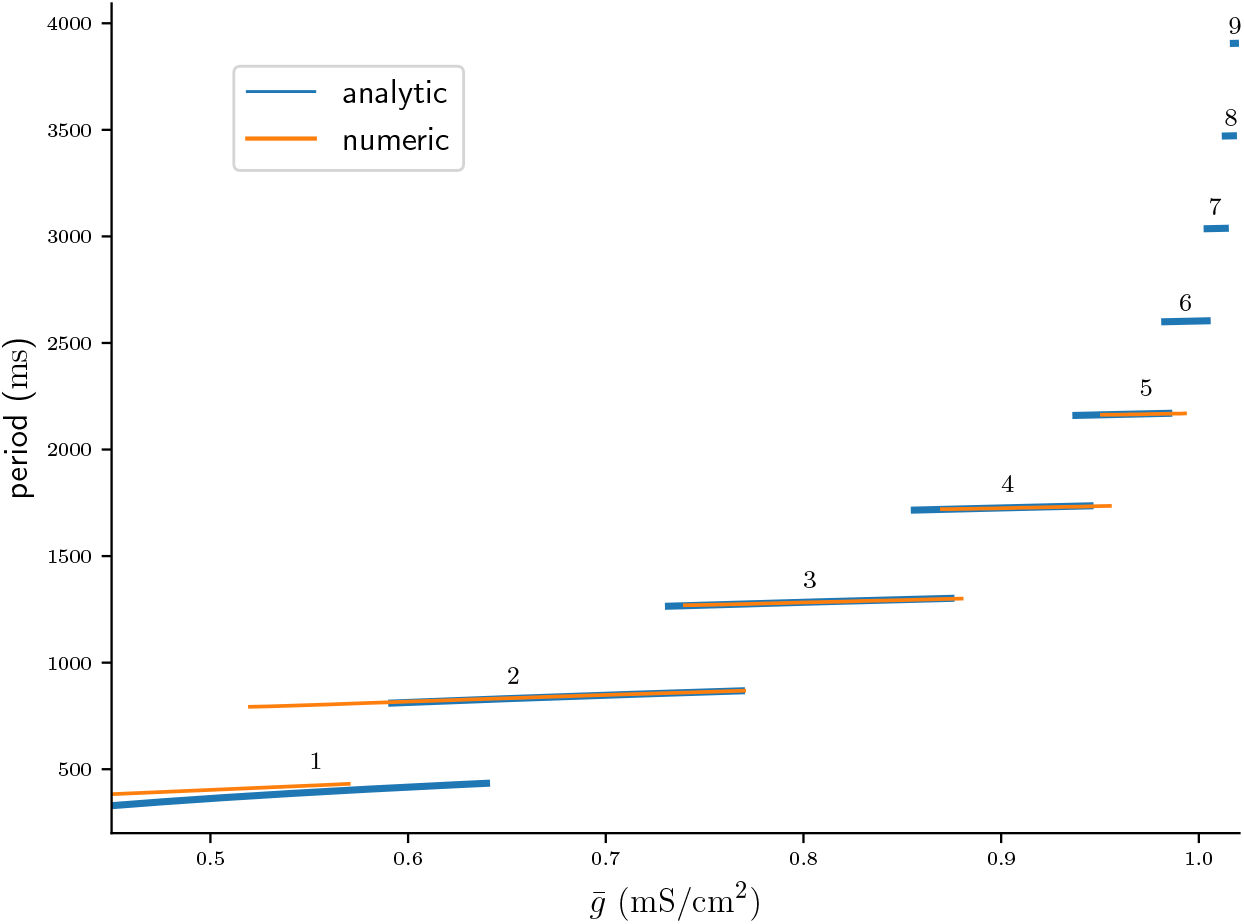
Bifurcation diagrams of stable *n* – *n* solutions computed analytically from fixed points of ∏_*n*_ and plotted on the respective intervals of 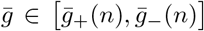 (blue), and computed from numerical integrations of the ODEs (orange).

As previously observed in fig. 11B our maps prediction of the cycle period is accurate for *n* > 1. Recall that our reduction assumptions required a sufficiently large coupling strength, which we numerically estimated to be 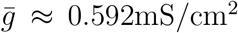 in fig. 6. The mismatch in period for 1 – 1 solutions, but also the mismatch in the left bifurcation border 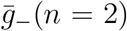 of the 2 – 2 solution can be attributed to the violation of that assumption. However, even for branches of large *n* – *n* solutions there is a mismatch between the bifurcation borders. Presumably our assumptions on the time scales of *w* and *s* dynamics do not hold here, and can only be captured by more complex approximations. Nevertheless, our map allows approximate extrapolation of the cycle period and the respective bifurcation borders where numerical integration of the ODEs would require a very small time step.

## 4. Discussion

Synaptic depression of inhibition is believed to play an important role in the generation of rhythmic activity involved in many motor rhythms such as in leech swimming (Mangan et al., 1994) and leech heart beat (Calabrese et al., 1995), and in the lobster pyloric system (Manor et al., 1997; Rabbah & Nadim, 2007). In half-centre CPGs synaptic depression can act as a burst termination mechanism, enabling the alternation of bursting between the two sides of the CPG (Brown, 1911). The struggling escape behaviour in *Xenopus* tadpoles is one such motor rhythm that is thought to be driven by synaptic depression of reciprocal inhibition between the two sides of the tadpole spinal cord (Li et al., 2007). Mathematical modelling can shed light on the underlying mechanisms that enable the generation of such anti-phase bursts in struggling, and help identify the components that control this rhythm allowing it to switch between different patterns.

To study the mechanisms of burst generation in tadpole struggling we have analysed a neuronal model network that consists of a pair of inhibitory neurons that undergo a frequency dependent synaptic depression. When the strength of synaptic inhibition between the neurons is varied, such a simple network can display a range of different struggling-like *n* – *n* burst patterns. Using the timescale disparity between neuronal and synaptic dynamics, we have reduced the network model of eight ODEs to a scalar first return map ∏_*n*_ of the slow depression variable *d*. This map ∏_*n*_ is a composition of two maps, *F_n_* and *Q_n_*, that model the evolution of the depression during the free and quiet phases of *n* – *n* solutions respectively. Both *F_n_* and *Q_n_* maps are constructed by using the dynamics of single uncoupled neurons. Fixed points of ∏_*n*_ are created in pairs through a fold bifurcation of maps, where the stable fixed point correspond to stable *n* – *n* burst solutions of the full two-cell system of ODEs. The results from our one-dimensional map match excellently with numerical simulation of the full struggling network. In line with Brown’s 1911 rhythmogenesis hypothesis, our results support the idea that synaptic depression in tadpoles may act as a burst termination mechanism which produces the anti-phase patterns found in struggling (Li et al., 2007).

We have studied *n* – *n* solutions assuming that the synaptic coupling 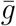 between the two cells is symmetrical. However, Bose & Booth (2011) have shown that asymmetrical coupling 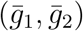 can result in network solutions of type *m* – *n*, where one cell fires *m* spikes, while the other *n* spikes. It is conceivable that our map construction can be extended to also capture such *m* – *n* solutions. Remember, in the case of symmetrical coupling with *n* – *n* solutions, the timecourse of the depression variables *d*_1_ and *d*_2_ were in anti-phase, and it was therefore sufficient to track only one of the two variables. To capture the full network dynamics in case of asymmetrical coupling one would also have to account for burst patterns of type *m* – *n*, where the solutions of the depression variables *d*_1_ and *d*_2_ are not simply time-shifted versions of each other. To do that, one could track the state of both variables by constructing a two-dimensional Poincaré map ∏(*d*_1_, *d*_2_). While geometrical interpretation of two-dimensional maps remains challenging, there exist a number of recent studies which have employed novel geometrical analysis methods to understand the dynamics of two-dimensional maps of small neuronal networks (Akcay et al., 2014, 2018; Liao et al., 2020). Generally speaking, our map construction approach is applicable to any small network, even with more than two neurons. As long as the network dynamics occur on separable timescales the main challenges to the map construction lie in identifying the slowest variables, and finding an appropriate, simplified description of their respective timecourses. In theory, the reduction approach can be also applied to neuronal systems with more than two timescales (e.g. see Kuehn, 2015).

Struggling in tadpoles is initiated by an increase in cIN firing frequency, which has been hypothesised to lead to stronger synaptic depression of commissural inhibition and result in the iconic anti-phase bursting (Li et al., 2007). It would therefore be interesting to study how varying the cell intrinsic firing period *T* could affect the network rhythm. While we have laid out the framework to perform such an investigation, due to the choice of neural model we have avoided varying *T*. Recall that *T* is a derived parameter in the Morris & Lecar (1981) model, and can therefore not be varied in isolation of other model parameters. This makes verifying any analytical results from our map analysis via numerical integration of the ODEs difficult. A more abstract model such as the quadratic integrate-and-fire model (Izhikevich, 2004) allows varying *T* independently of other model parameters, and could be more fitting in this scenario.

Our simulations of the network showed that *n* – *n* solutions lose robustness as their period is increased. That is, solutions with a larger cycle period occur on increasingly smaller intervals of the coupling strength. We were able to replicate this finding by numerically finding the respective left and right borders of stable *n* – *n* branches of fixed points of ∏_*n*_, and showing that the distance between these borders shrinks with *n*. We have also noted the resemblance of our bifurcation diagram to one where such *n* – *n* branches are created via a period-increment bifurcation with co-existent attractors of scalar linear maps with a discontinuity (Avrutin et al., 2012, 2011). It is worthwhile noting that the bifurcations of piecewise linear maps studied by Avrutin et al. and colleagues result from a “reinjection” mechanism, first described by Perez (1985). Here the orbit of a map performs multiple iterations on one side of the discontinuity, before jumping to the other side and being *reinjected* back into the initial side of the discontinuity. The stark difference of such a map to our map is that reinjection allows a *single* scalar map to produce periodic solutions of varying periods. In contrast, we rely on *n* different maps ∏_*n*_ to describe the dynamics burst dynamics without explicitly capturing the period increment bifurcations. It is therefore conceivable that despite the complexity and non-linearity of the dynamics of our two-cell network, a single piecewise-linear map might be already sufficient to capture the mechanisms that shape the parameter space of the full system. In their discussion, Bose & Booth (2011) briefly outline ideas about how such a linear map could be constructed.

In addition to stable *n* – *n* solutions, the numerical continuation by Bose & Booth (2011) also revealed branches of unstable *n* – *n* solutions. While we have identified fold bifurcations of our burst map, we have not found corresponding bifurcations of the flow ODE system, and have generally ignored the significance of unstable map fixed points. However, our results are reminiscent of a saddle homoclinic orbit bifurcation, where a stable periodic orbit collides with a saddle equilibrium, and its period becomes infinite. A number of theoretical studies have investigated such a *homoclinic tangency to a periodic orbit* in higher-dimensional systems (e.g. Gavrilov & Shilnikov, 1972; Gaspard & Wang, 1987), and a next step would be to provide a rigorous explanation of not only the map dynamics, but also of the flow dynamics of the ODE system.

We have shown that when the strength of the maximum synaptic conductance is varied, synaptic depression of inhibition can enable our two-cell network to produce burst solutions of different periods. This result is in line with the idea that one role of synaptic depression in the nervous system may be to allow a finite size neuronal network to participate in different tasks by producing a large number of rhythms (Bose & Booth, 2011; Jalil et al., 2004; Li et al., 2007). To change from one rhythm to another would only require a reconfiguration of the network through changes in synaptic coupling strength, which can occur through the process of learning. Thus short-term synaptic depression of inhibition may provide means for a network to adapt to environmental challenges without changing its topology, that is without the introduction or removal of neurons.

## 5. Acknowledgements

## Funding

This work was supported by the Wellcome Trust Doctoral Training Programme in Neural Dynamics, Grant no. 099699/Z/12/Z.

We thank Alan Champneys for sharing his insights on non-continuous maps during the course of this research. We are also grateful to Alan Roberts and Stephen R. Soffe for their comments on earlier versions of the manuscript.

## Appendix A. Model equations and parameters

The asymptotic functions *m*_∞_ and *w*_∞_ for the calcium and potassium conductances, respectively, are given by

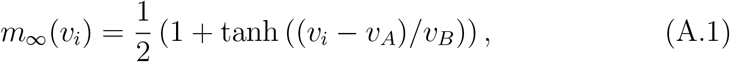

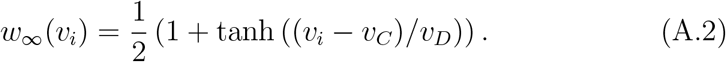

Model parameters were adapted from Bose & Booth (2011) and are given in table A.1:

## Appendix B. Computing bifurcation diagram numerically

The bifurcation diagram of stable *n*–*n* solutions of the two-cell network in fig. 4 is obtained numerically as follows: We initialise the coupling strength at parameter values associated with one type of *n* – *n* solution, that is we choose the values 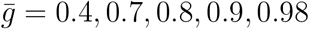 for the 1 – 1, 2 – 2, 3 – 3, 4 – 4, and 5 – 5 solutions respectively. For each 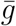 the system is then numerically integrated sufficiently long for any transients to fully subside. We then identify one period of the solution by finding the first return of the depression variable *d*_1_. That is, we choose some value *d_k_* at a spike time *t_k_*, and by iterating from spike to spike find some subsequent value *d*_*k*+1_ at spike time such that |*d*_*k*+1_ – *d_k_* | < *ϵ*. If a periodic solution of type *n* – *n* is found in such way, 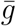 is step-wise increased/decreased, and the above algorithm is repeated. Otherwise, the set of all previously found solutions and the corresponding values 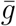 are returned.

**Table A.1:** Default parameters for coupled Morris-Lecar model.

| Parameter | value |
| --- | --- |
| $g_L$ | 0.15 mS/cm<sup>2</sup> |
| $g_{Ca}$ | 0.3 mS/cm<sup>2</sup> |
| $g_K$ | 0.6 mS/cm<sup>2</sup> |
| $v_L$ | -50 mV |
| $v_{Ca}$ | 100 mV |
| $v_K$ | -70 mV |
| $v_A$ | 1 mV |
| $v_B$ | 14.5 mV |
| $v_C$ | 4 mV |
| $v_D$ | 15 mV |
| $I$ | 6 μA/cm<sup>2</sup> |
| $\tau_w$ | 100 ms |
| $\tau_d$ | 1000 ms |
| $\tau_s$ | 100 ms |
| $v_\theta$ | 0 mV |
| $v_s$ | -80 mV |
| $\lambda$ | 0.5 |
| $T$ | 217 ms |
| $g^*$ | 0.03804 mS/cm<sup>2</sup> |

Note that applying conventional methods of numerical continuation is not straightforward for our model, as the discontinuous resets in eq. (7) and (8) make the root finding challenging (e.g. see Kuznetsov, 2004, for continuation methods).

## References

Akcay, Z., Bose, A., & Nadim, F. (2014). Effects of synaptic plasticity on phase and period locking in a network of two oscillatory neurons. The Journal of Mathematical Neuroscience, 4, 8. doi:10.1186/2190-8567-4-8.

Akcay, Z., Huang, X., Nadim, F., & Bose, A. (2018). Phase-locking and bistability in neuronal networks with synaptic depression. Physica D: Nonlinear Phenomena, 364, 8 – 21. doi:10.1016/j.physd.2017.09.007.

Avrutin, V., Granados, A., & Schanz, M. (2011). Sufficient conditions for a period incrementing big bang bifurcation in one-dimensional maps. Non-linearity, 24, 2575–2598. URL: https://doi.org/10.1088/0951-7715/24/9/012. doi:10.1088/0951-7715/24/9/012.

Avrutin, V., Schanz, M., & Schenke, B. (2012). Breaking the continuity of a piecewise linear map. ESAIM: Proceedings, 36, 73–105. URL: https://doi.org/10.1051/proc/201236008. doi:10.1051/proc/201236008.

Bose, A., & Booth, V. (2011). Co-existent activity patterns in inhibitory neuronal networks with short-term synaptic depression. Journal of Theoretical Biology, 272, 42–54. doi:10.1016/j.jtbi.2010.12.001.

Bose, A., Manor, Y., & Nadim, F. (2001). Bistable oscillations arising from synaptic depression. SIAM Journal on Applied Mathematics, 62, 706–727. doi:10.1137/S0036139900378050.

Brown, T. G. (1911). The intrinsic factors in the act of progression in the mammal. Proceedings of the Royal Society of London. Series B, 84, 308–319.

Calabrese, R. L., Nadim, F., & Olsen, Ø. H. (1995). Heartbeat control in the medicinal leech: A model system for understanding the origin, coordination, and modulation of rhythmic motor patterns. Journal of neurobiology, 27, 390–402. doi:10.1002/neu.480270311.

Clarke, J. D., Hayes, B. P., Hunt, S. P., & Roberts, A. (1984). Sensory physiology, anatomy and immunohistochemistry of Rohon-Beard neurones in embryos of *Xenopus laevis*. The Journal of Physiology, 348, 511–525. doi:10.1113/jphysiol.1984.sp015122.

Donovan, M., Wenner, P., Chub, N., Tabak, J., & Rinzel, J. (1998). Mechanisms of spontaneous activity in the developing spinal cord and their relevance to locomotion. Annals of the New York Academy of Sciences, 860, 130–141.

Ermentrout, B. (2002). Simulating, analyzing, and animating dynamical systems: a guide to xppaut for researchers and students. SIAM.

Ermentrout, G. B., & Terman, D. H. (2010). Mathematical foundations of neuroscience volume 35 of Interdisciplinary Applied Mathematics. Springer New York. doi: 10.1007/978-0-387-87708-2.

Friesen, W. O. (1994). Reciprocal inhibition: a mechanism underlying oscillatory animal movements. Neuroscience & Biobehavioral Reviews, 18, 547–553. doi:10.1016/0149-7634(94)90010-8.

Gardini, L., Avrutin, V., Schanz, M., Granados, A., & Sushko, I. (2012). Organizing centers in parameter space of discontinuous 1d maps. the case of increasing/decreasing branches. ESAIM: Proceedings, 36, 106–120. URL: https://doi.org/10.1051/proc/201236009. doi:10.1051/proc/201236009.

Gaspard, P., & Wang, X. J. (1987). Homoclinic orbits and mixed-mode oscillations in far-from-equilibrium systems. Journal of Statistical Physics, 48, 151–199. doi:10.1007/BF01010405.

Gavrilov, N. K., & Shilnikov, L. P. (1972). On three-dimensional dynamical systems close to systems with a structurally unstable homoclinic curve. I. Mathematics of the USSR-Sbornik, 17, 467. doi:10.1070/SM1972v017n04ABEH001597.

Gonzalez, A. S. (2021). pynverse. https://github.com/alvarosg/pynverse.

Izhikevich, E. M. (2004). Which model to use for cortical spiking neurons? ieee transactions on neural networks, 15, 1063–1070. doi:10.1109/TNN.2004.832719.

Jalil, S., Grigull, J., & Skinner, F. K. (2004). Novel bursting patterns emerging from model inhibitory networks with synaptic depression. Journal of Computational Neuroscience, 17, 31–45. doi:10.1023/B:JCNS.0000023870.23322.0a.

Kahn, J. A., & Roberts, A. (1982). The neuromuscular basis of rhythmic struggling movements in embryos of *Xenopus laevis*. Journal of Experimental Biology, 99, 197–205.

Kahn, J. A., Roberts, A., & Kashin, S. (1982). The neuromuscular basis of swimming movements in embryos of the amphibian *Xenopus laevis*. Journal of Experimental Biology, 99, 175–184.

Kuehn, C. (2015). *Multiple Time Scale Dynamics* volume 191 of *Applied Mathematical Sciences*. Springer International Publishing. doi:10.1007/978-3-319-12316-5.

Kuznetsov, A. (2004). Elements of applied bifurcation theory. Springer Science & Business Media. doi:10.1007/978-1-4757-3978-7.

Li, W.-C., Sautois, B., Roberts, A., & Soffe, S. R. (2007). Reconfiguration of a vertebrate motor network: specific neuron recruitment and context-dependent synaptic plasticity. The Journal of Neuroscience, 27, 12267–12276. doi:10.1523/JNEUROSCI.3694-07.2007.

Liao, G., Diekman, C., & Bose, A. (2020). Entrainment Dynamics of Forced Hierarchical Circadian Systems Revealed by 2-Dimensional Maps. SIAM J. Appl. Dyn. Syst.,. URL: https://epubs.siam.org/doi/abs/10.1137/19M1307676.

Mangan, P., Cometa, A., & Friesen, W. (1994). Modulation of swimming behavior in the medicinal leech. IV. Serotonin-induced alteration of synaptic interactions between neurons of the swim circuit. Journal of comparative physiology. A, Sensory, neural, and behavioral physiology, 175, 723–736. doi:10.1007/BF00191844.

Manor, Y., Nadim, F., Abbott, L., & Marder, E. (1997). Temporal dynamics of graded synaptic transmission in the lobster stomatogastric ganglion. Journal of Neuroscience, 17, 5610–5621. doi:10.1523/JNEUROSCI.17-14-05610.1997.

Matveev, V., Bose, A., & Nadim, F. (2007). Capturing the bursting dynamics of a two-cell inhibitory network using a one-dimensional map. Journal of Computational Neuroscience, 23, 169–187. doi:10.1007/s10827-007-0026-x.

Morris, C., & Lecar, H. (1981). Voltage oscillations in the barnacle giant muscle fiber. Biophysical Journal, 35, 193–213. doi:10.1016/S0006-3495(81)84782-0.

Nadim, F., & Manor, Y. (2000). The role of short-term synaptic dynamics in motor control. Current Opinion in Neurobiology, 10, 683–690. doi:10.1016/S0959-4388(00)00159-8.

Nadim, F., Manor, Y., Kopell, N., & Marder, E. (1999). Synaptic depression creates a switch that controls the frequency of an oscillatory circuit. Proceedings of the National Academy of Sciences, 96, 8206–8211. doi:10.1073/pnas.96.14.8206.

Olenik, M. (2021). PyXPP. https://github.com/markolenik/PyXPP.

Perez, J. M. (1985). Mechanism for global features of chaos in a driven nonlinear oscillator. Physical Review A, 32, 2513–2516. URL: https://doi.org/10.1103/physreva.32.2513. doi:10.1103/physreva.32.2513.

Perkel, D. H., & Mulloney, B. (1974). Motor pattern production in reciprocally inhibitory neurons exhibiting postinhibitory rebound. Science, 185, 181–183. doi:10.1126/science.185.4146.181.

Rabbah, P., & Nadim, F. (2007). Distinct synaptic dynamics of heterogeneous pacemaker neurons in an oscillatory network. Journal of neurophysiology, 97, 2239–2253. doi:10.1152/jn.01161.2006.

Reiss, R. F. (1962). A theory and simulation of rhythmic behavior due to reciprocal inhibition in small nerve nets. In Proceedings of the May 1-3, 1962, spring joint computer conference (pp. 171–194). ACM. doi: 10.1145/1460833.1460854.

Soffe, S. R. (1993). Two distinct rhythmic motor patterns are driven by common premotor and motor neurons in a simple vertebrate spinal cord. The Journal of Neuroscience, 13, 4456–4469.

Tramontana, F., Gardini, L., Avrutin, V., & Schanz, M. (2012). Period adding in piecewise linear maps with two discontinuities. International Journal of Bifurcation and Chaos, 22, 1250068. URL: https://doi.org/10.1142/s021812741250068x. doi:10.1142/s021812741250068x.

Tsodyks, M. V., & Markram, H. (1997). The neural code between neo-cortical pyramidal neurons depends on neurotransmitter release probability. Proceedings of the National Academy of Sciences, 94, 719–723. doi:10.1073/pnas.94.2.719.

Virtanen, P., Gommers, R., Oliphant, T. E., Haberland, M., Reddy, T., Cournapeau, D., Burovski, E., Peterson, P., Weckesser, W., Bright, J., van der Walt, S. J., Brett, M., Wilson, J., Millman, K. J., Mayorov, N., Nelson, A. R. J., Jones, E., Kern, R., Larson, E., Carey, C. J., Polat, İ., Feng, Y., Moore, E. W., VanderPlas, J., Laxalde, D., Perktold, J., Cimrman, R., Henriksen, I., Quintero, E. A., Harris, C. R., Archibald, A. M., Ribeiro, A. H., Pedregosa, F., van Mulbregt, P., & SciPy 1.0 Contributors (2020). SciPy 1.0: Fundamental Algorithms for Scientific Computing in Python. Nature Methods, 17, 261–272. doi:10.1038/s41592-019-0686-2.

